# An Oscillatory Network Model of Head Direction, Spatially Periodic Cells and Place Cells Using Locomotor Inputs

**DOI:** 10.1101/080267

**Authors:** Karthik Soman, Vignesh Muralidharan, V. Srinivasa Chakravarthy

## Abstract

We propose a computational modeling approach that explains the formation of a range of spatial cells like head direction cells, grid cells, border cells and place cells which are believed to play a pivotal role in the spatial navigation of an animal. Most existing models insert special symmetry conditions in the models in order to obtain such symmetries in the outcome; our models do not require such symmetry assumptions. Our modeling approach is embodied in two models: a simple one (Model #1) and a more detailed version (Model #2). In Model #1, velocity input is presented to a layer of Head Direction cells, with no special topology requirements, the outputs of which are presented to a layer of Path Integration neurons. A variety of spatially periodic responses resembling grid cells, are obtained using the Principal Components of Path Integration layer. In Model #2, the input consists of the locomotor rhythms from the four legs of a virtual animal. These rhythms are integrated into the phases of a layer of oscillatory neurons, whose outputs drive a layer of Head Direction cells. The Head Direction cells in turn drive a layer of Path Integration neurons, which in turn project to two successive layers of Lateral Anti Hebbian Networks (LAHN). Cells in the first LAHN resemble grid cells (with both hexagonal and square gridness), and border cells. Cells in the second LAHN exhibit place cell behaviour and a new cell type known as corner cell. Both grid cells and place cells exhibit phase precession in 1D and 2D spaces. The models outline the neural hierarchy necessary to obtain the complete range of spatial cell responses found in the hippocampal system.

## Introduction

The discovery of place cells, cells that fire whenever the animal visits a certain location in the ambient space, in the CA1 field of hippocampus, paved the way to the study of how space is represented in the temporal lobe structures, and further led to the discovery of a larger class of cells called ‘spatial cells’(Hafting et al., 2005; O’Keefe and Dostrovsky, 1971; Solstad et al., 2008; Taube et al., 1990a; Taube et al., 1990b)

Place cells give an accurate representation of the current location of the animal in space. But this alone is not sufficient for efficient navigation of an animal. There must be a dedicated neuronal system to compute the sense of direction and distance that the animal traverses. Ranck *et al* (1990) discovered a group of neurons from the postsubiculum region that fired only when the animal’s head is in a particular direction in the horizontal plane (yaw plane) (Taube et al., 1990a; Taube et al., 1990b). The activity of these head direction cells is independent of the place and the behavior of the animal and the direction in which they fire is known as the “preferred direction” of the cell. Hence head direction cells are regarded as the “internal compass” that gives a sense of direction(Valerio and Taube, 2012).

Place cells are mainly found in the CA1 region of the hippocampus. The conundrum of what formed input to the place cell activity remained elusive until it was shown that place cell activity persisted even after the lesion of all intra-hippocampal inputs to CA1 subfield indicating that the input for place cell activity must be coming from outside the hippocampus (Brun et al., 2002).Studies showed that the dorsal medial entorhinal cortex (mEC) was a prime candidate because of its vast connections to the CA1 subfield. Edvard Moser, May Britt Moser and colleagues(Hafting et al., 2005) delineated a group of neurons in mEC that had a very striking firing field with a regular organization such that the multiple firing fields of a single neuron roughly formed the vertices of a hexagon. The firing field thus tessellates the space that the animal forages and forms a hexagonal grid-like pattern and hence the name grid cells. Further recording from other regions of mEC proved that grid fields varied in certain characteristics such as grid scale and grid width. Grid scale and grid width progressively increase on moving from dorsal to ventral region of mEC (Brun et al., 2008; Stensola et al., 2012). Grid cells are assumed to perform the function of path integration. Grid cells are more stable to the removal of external stationary cues like visual landmarks or olfactory cues (Moser et al., 2014). Hence in this study we model the formation of grid cells using proprioceptive inputs (locomotion)and the evolution of firing field in the model.

Moser *et al* also reported neurons from mEC which fired when the animal was close to the borders of the environment (Solstad et al., 2008).These border cells firing persisted even after manipulating the environment by stretching and also in environments of different shapes and sizes. Border cells or Boundary Vector Cells (BVC) were also reported to be located in the subiculum region of the hippocampal formation(Lever et al., 2009).The aforementioned cells such as head direction cells, grid cells, border cells and place cells seem to be the building blocks of a comprehensive neuronal network for representing and negotiating space.

In this study, we present a model of several types of hippocampal spatial cells including grid cells, place cells, head direction cells and border cells. Existing models of grid cells fall into two broad categories: oscillatory interference models and attractor network models. The Oscillatory Interference model was originally proposed by O’Keefe and Recce(O’Keefe and Recce, 1993) to explain the form of temporal coding of place cells known as *phase precession*. As the animal moves through the place field, the phase of the place cell spiking activity precesses with respect to a constant background theta rhythm. The essence of this model is that the interference between two sub-threshold membrane potential oscillations (MPO) with slightly different frequencies can result in an interference pattern such that a proper thresholding will give rise to spiking over spatially periodic locations. Frequency of one MPO is kept constant while the other is modulated by the speed and direction of motion (Burgess et al., 2007). Albeit this model explains phase precession of a place cell, it is not able to account for the periodic spiking which seems to be unreliable for a place cell having a localized firing field. The model seems to fit quite well for the grid cell because of its spatially periodic firing field. Burgess*et al* proposedan interference model (Burgess et al., 2007), similar to that of (O’Keefe and Recce, 1993), but generalized it to explain the grid field formation in two dimensional space. The model assumes that subthreshold MPOs of multiple dendrites of a grid cell in medial entorhinal cortex (mEC) are modulated by the velocity of the animal such that the velocity is projected to those dendritic directions which are multiples of 60^°^. In other words, the model is based on an unrealistic constraint that the preferred head directions are sharply concentrated around integral multiples of 60^°^. Each modulated dendritic MPO was assumed to interfere with a subthreshold unmodulated baseline MPO. Further thresholding and remapping gave rise to perfect hexagonal firing fields. Many variations of this model have been introduced (Blair et al., 2008; Hasselmo, 2008; Zilli and Hasselmo, 2010) thereafter but the basic principle remains the same. The merit of this model is that the resetting of path integration takes place naturally because of the inherent periodicity in the oscillations rather than using hard resets like modulo functions(Gaussier et al., 2007). But the model is biologically unrealistic because of the 60^°^ constraint on the head direction system; it constrains the number of head direction cells that can be implemented in the model.

Grid cell models based on continuous attractor neural network models (Burak and Fiete, 2009; Fuhs and Touretzky, 2006; Guanella et al., 2007; McNaughton et al., 2006) consist of 2D layers of neurons in which, each neuron has circularly symmetric ON-center, OFF-surround lateral connectivity (Burak and Fiete, 2009; Fuhs and Touretzky, 2006). Such models exhibit hexagonal pattern formation in the neural space by Turing instability(Turing, 1952). To simulate the empirical grid firing field, the animal’s motion is coupled to the network such that the hexagonal pattern formed gets translated on the neural space based on the animal’s velocity. This path integration is achieved in the network by supposing the existence of dynamic, velocity-modulated synaptic weights,such as an asymmetric componentwhich is proportional to the velocity. This asymmetric component is added to the centre surround symmetric connectivity of each neuron. This allows the velocity of the animal to be coupled to the network dynamics, resulting in translation of the formed pattern on the neural space. Probing a single neuron from the neural network and remapping its response onto the external space produces a hexagonal firing field. This type of neural architecture assumes a toroidal topology to avoid the edge effect (Guanella et al., 2007; McNaughton et al., 2006). A similar approach has been used to model head direction cells also. Typical models of head direction cell network (Knierim et al., 1995; Redish et al., 1996; Sharp et al., 2001; Zhang, 1996) assume a ring topology with connections following the same aforementioned pattern. The network’s activity bump moves when the animal turns, which renders the lateral connections asymmetric. Thus the model offers a computational mechanism for rotational path integration. For the shift between symmetric to asymmetric connections to be stable, Zang*et al* proposed a spatial derivative law such that the asymmetry component must be proportional to the spatial derivative of the symmetry component of the weight connection(Zhang, 1996).

From the aforementioned corpus of literature we see that there is a need for a comprehensive model that can incorporate the formation of all the spatial cells in a biologically realistic self-organized fashion. Here we propose a model of spatial cell formation which attempts to circumvent the shortcomings of the aforementioned models.

## 2.0 Methods

In this section, we present two models of hippocampal spatial cells: Model 1 and Model 2. Model 1 is a simpler version of Model 2, but is more transparent due to its simplicity. It reveals a key insight in our modeling approach. Model 1explains how periodicity arises in the spatial cell responses though there is no periodicity in the input, nor is there any special symmetry present in the network architecture. Some of the simplifying modeling assumptions of Model 1are then relaxed, thereby producing Model 2.

The models have certain common architectural elements. In both, motion-related inputs are presented to a layer of Head Direction (HD) cells, which is a layer of neurons without any special symmetries (Taube et al., 1990a; Taube et al., 1990b). The HD layer responses are integrated in the next layer known as the Path Integration (PI) layer. The PI layer in turn projects, via trainable connections, to another layer where a variety of spatial cells, particularly a variety of grid cells, naturally emerge. This is termed the Spatial Cell (SC) layer. Model 2 has an additional stage: the outputs of the SC layer project to another layer, the Place Cell (PC) layer, where place cell responses emerge.

A virtual animal is made to forage inside a square box of size 3 units. Trajectories of the animal, that involve an upper limit on curvature, are constructed using a method described in Appendix A1.1.In Model 1, the virtual animal is represented as a point, and its motion is represented explicitly in terms of speed, s, and direction, θ. In Model 2, the virtual animal is represented as a four-legged creature and motion is represented in terms of four locomotor rhythms generated by the legs.

### 2.1 Model 1

The proposed model has threelayers, HD layer,Path Integration (PI) layer and an output SC layerthat represents the region of Entorhinal Cortex (EC) to which the PIresponseconverges as input (Fig. 1). The response of i^th^ HDcell is computed as the projection of the animal’s current direction on to the i^th^ preferred direction, given as,

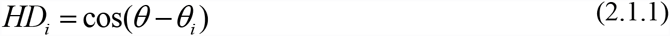

**Fig. 1:**
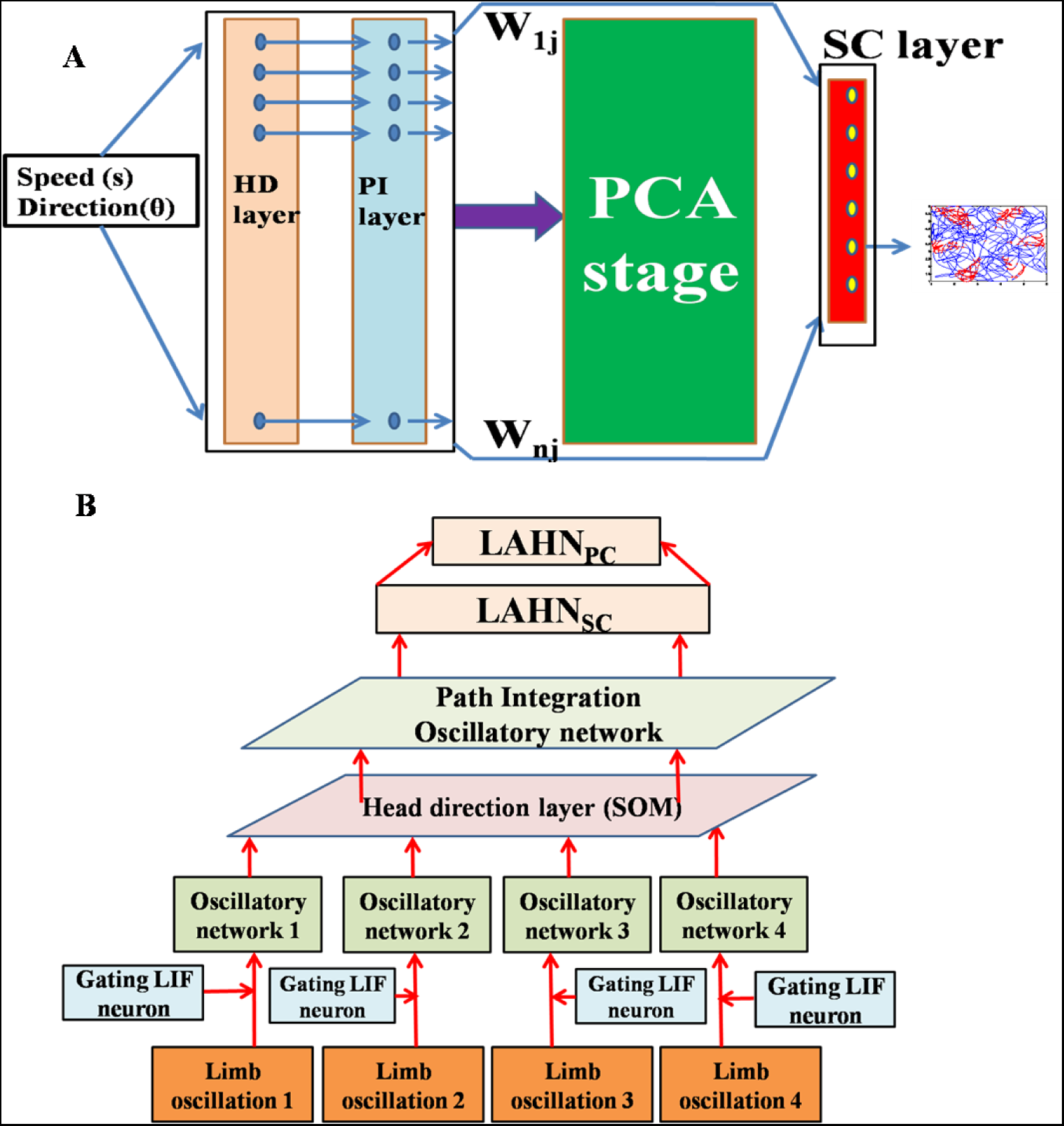
The architecturesfor (A) Model 1, (B) Model 2.

θ is the current heading direction and θ_i_ is the preferred direction of i^th^ HD cell.

The outputs of HD layer project to the subsequent PI layer, via one-to-one connections of unity strength. PI neurons are modelled as oscillatory neurons whose phases encode the result of path integration performed on HD cell responses. Each HD cell modulates the corresponding PI neuron’s dendritic frequency from its base frequency *f*_0_ thereby changing the phase of the PI neuron. The response of the i^th^ PI cell is given as,

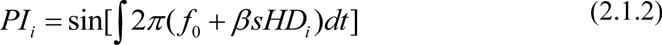

– *β* is a spatial scaling parameter
– s is speed of the animal

The PI cell responses of i^th^ PI neuron, (PI_i_) are then thresholded by the following rule,

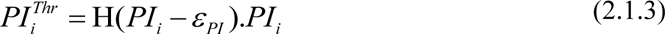

where, H is the Heaviside function, and *ε*_*PI*_ is the threshold value.

The thresholded PI values are projected via a linear weight stage (W^PC^) to a subsequent layer, the SC layer. Weight (W_ij_^PC^) from i^th^ PI^Thr^ to j^th^ SC layer neuronis computed by performing Principal Component Analysis (PCA) over PI^Thr^. The response of i^th^ neuron in the SC layer is given as

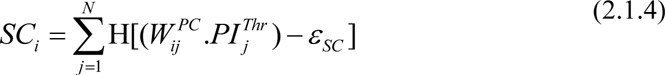

where, H is Heaviside function,

N is the number of PI neurons,

*ε*_*SC*_ is the threshold value.

The top few components of the computed principal components (PC) will be shown to reveal a variety of spatial cell-like responses including grid cells (both hexagonal and square grid cells), and cornercells (whose firing fields are at the corners of the space) as shown in results section. The emergence of spatially periodic firing field is due to the inherent periodicity in the PC weights. The neurons that receive PCs whose peaks are separated by ≈ 60^°^(PC 6 and PC 7) show hexagonal grid cell like activity (See appendix 2).The firing field of each neuron is depicted by placingred dots on those positions of the trajectory where the respective neuron in the SC layeris active (Fig 3.A.1-3.F.1).

**Fig 3:**
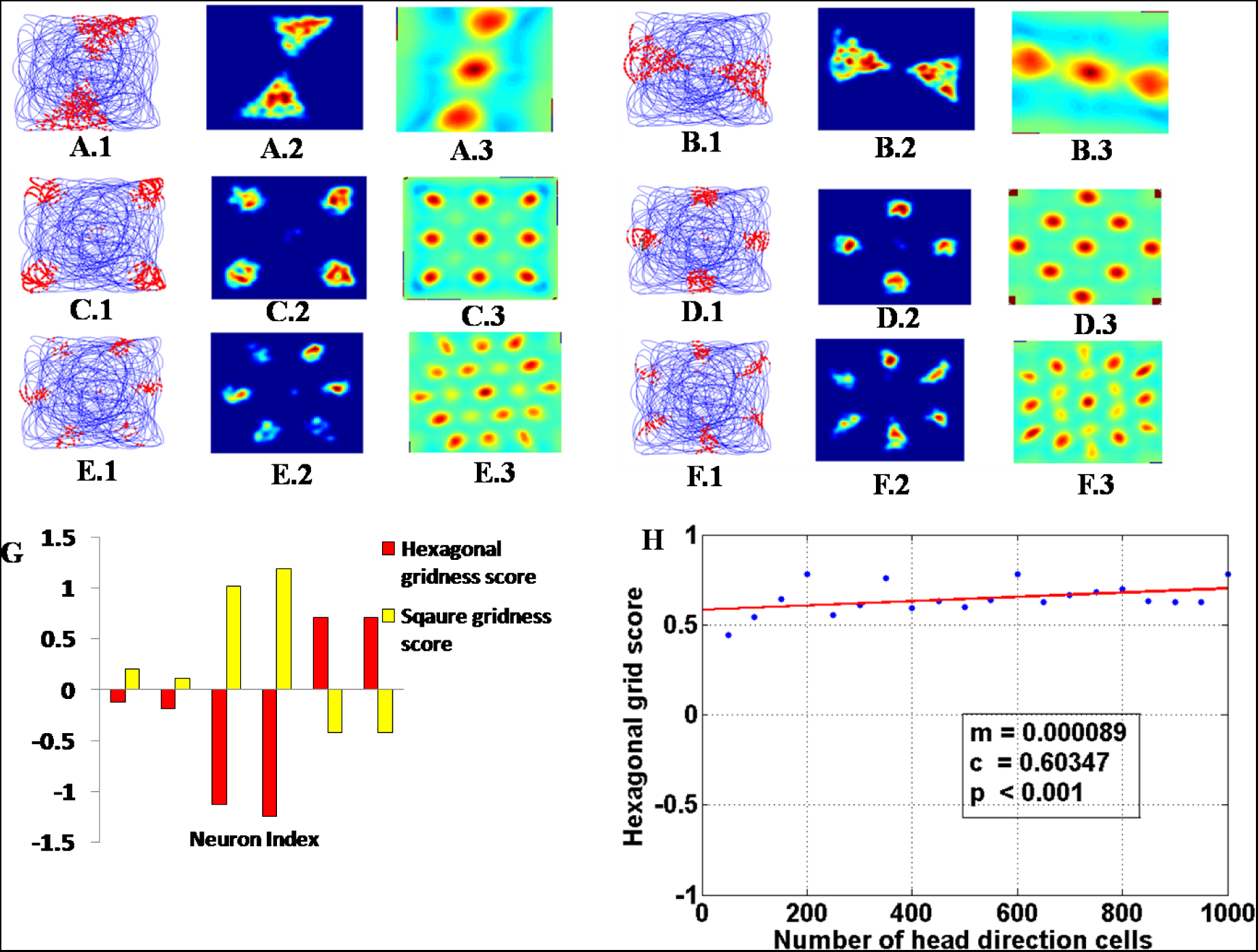
(A-F)The firing fields (A.1-F.1), rate map (A.2-F.2) and the autocorrelation maps (A.3-F.3) from the SC layer in the analytical model. (G)The gridness scores (hexagon and square) of the SC layer neurons.(H) The regression analysis between hexagonal gridness score and number of head direction cells shows the independence of the hexagonal grid formation to the number of head direction cells.

### 2.2. Model 2

Some of the simplifying assumptions of Model 1 are relaxed to yield a biologically more plausible Model 2. A novel element in Model 2 is the manner in which motion-related input is represented. A majority of models of spatial navigation that incorporate path integration, explicitly assume the coordinates x and y, or the velocity v_x_ and v_y_, of the animal. But such an assumption is unrealistic since the animal’s nervous system does not have access to such an explicit representation of its position or velocity in an arbitrary coordinate system. These motion-related quantities are inferred by the nervous system from thesensory streams – visual, proprioceptive etc. In Model 2, we consider the proprioceptive locomotor input as a source of motion-related information.

In this model (Fig 1B), the animal is represented, not as a point mass, but as a four-legged creature, with its four limbs located at the corners of a roughly rectangular strip (FigA3.A inAppendix A3). The center of mass of the animal, which is located at the center of the rectangular strip formed by the four legs, moves along a synthesized trajectory at a speed, s. The trajectory of the animal cuts along the midline of the rectangular strip.

The motion of each limb of the four-legged virtual animal is thought to be represented by a single oscillator defined by a unique phase. Oscillations corresponding to the four legs have the same oscillatory frequency. The details of how the four limb oscillations are generated as the virtual animal moves along a synthetic trajectory at a given speed, s, are described in Appendix A3. The limb oscillations are described as,

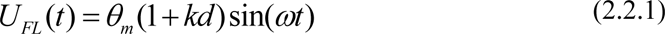

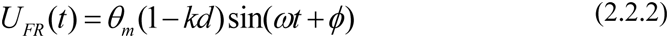

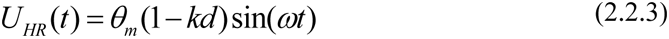

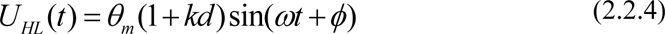

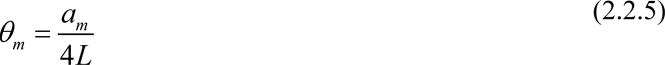

*U*_*FR*_ and*U*_*FL*_ = Fore-right and fore-left limb oscillations respectively.

*U*_*HR*_ and*U*_*HL*_ = Hind-right and hind-left limb oscillations respectively.

θ_m_= Angle made by the animal’slimb with respect to the normal line passing through the center of mass.

*a*_*m*_ = distance travelled by the center of mass of the animal while the limb completes one full cycle.

L = Length of the animal’s leg.

*φ* = Phase difference between the limb oscillations

*d* = spacing between the limb pair in the front (hind)

*k* = curvature of the trajectory

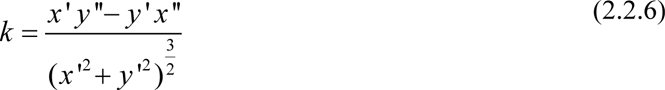

Eqn. (2.2.6) is the expression for the curvature for parametrically defined curves, where x and y denote the animal’s position on the trajectory. Note that in eqns. (2.2.1-2.2.4) above, the fore-right (fore-left) and hind-left (hind-right) rhythms are assumed to be in phase. Such a locomotor rhythm is defined as trot rhythm, in animal motion literature(Collins and Richmond, 1994).

Thelimb oscillations of eqn.2.2.1-4 are gated using a Leaky Integrate and Fire (LIF) neuron that spikes at a fixed frequency so that the oscillatory proprioceptive inputs are converted to proprioceptive pulses. The LIF neuron is modeled as:

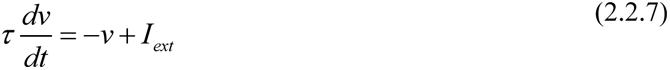

τ is time constant of the LIF neuron

v is the membrane potential of the neuron

v_thresh_ is spiking threshold of the LIF neuron

I_ext_ is the external current applied to the neuron. The LIF neuron receives sinusoidal current having the same frequency as that of the limb oscillations.

S is the output spike from the LIF neuron.

v_reset_ is the resetting voltage of the LIF neuron.

Proprioceptive pulses are given as:

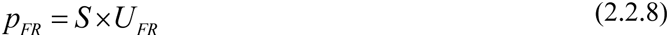

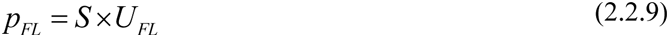

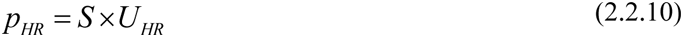

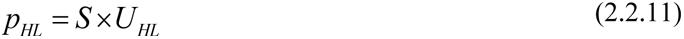

p_FR_,p_FL_,p_HR_and p_HL_are proprioceptive pulses from the respective limbs.

The proprioceptive pulses are passed onwards to four separate pools of oscillatory neuronal networks, O_FR_,O_FL_,O_HR_and O_HL_modeled as Kuramoto oscillators(Kuramoto, 1984). In each pool, the oscillatory neurons are coupled to each other by a coupling factor K. The response of O_FR_ oscillatory neuron is:

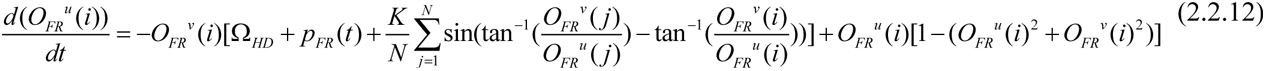

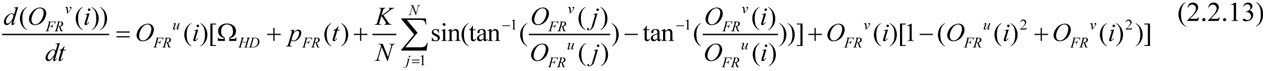

O_FR_^u^ and O_FR_^v^ are the u and v components of the phase of the oscillator O_FR_.

Ω_HD_is the base frequency,

K is the coupling factor,

N is the number of neurons in one cluster of oscillators.

(Note: Equations for dynamics of the responses of the remaining neurons, O_FL_, O_HR_ and O_HL_, can be written analogous to equations (2.2.12) and (2.2.13) simply by changing the subscript.)

Apart from the four oscillators (whose phases are modulated bythe proprioceptive pulses from the respective limbs) there exists a background oscillator (O_BG_) with the same base frequency as that ofthe other four oscillators butwhose phase remains independent of the proprioceptive pulses. We define a four-dimensional vector *Ψ*(*t*), as the vector of the instantaneous states of the four oscillators when the phase of O_BG_ reaches >π/2. Values of *Ψ*(*t*) are used to train a self-organizing map (SOM) of size 10x10, using the standard SOM algorithm (Kohonen, 1982).This SOM layer is analogous to the HD layer of Model 1.

Contrary to existing approaches (Knierim et al., 1995; Redish et al., 1996; Sharp et al., 2001; Zhang, 1996) to modeling head direction, that use a biologically unrealistic ring of neurons forming an attractor network, we use a more natural two-dimensional layer of neuronsto represent HD layer.

Equation (2.2.14) gives the response of the HDneuron.

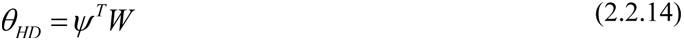

*Ψ*(*t*) is the normalized input state vector from the four oscillators. The weight vector of each neuron in the HD layer is also normalized. Therefore,the response of HD neuron is the cosine of the angle between *Ψ*(*t*) and W vectors since both vectors are normalized.

θ_HD_ is passed onward to a 10x10 arrayof PI oscillatory neuron layer which has one-to-one connections with the HD neuron layer. The phase of the neuron at location (i, j) in the PI layer is computed using (2.2.15) and (2.2.16).

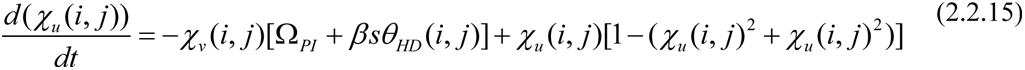

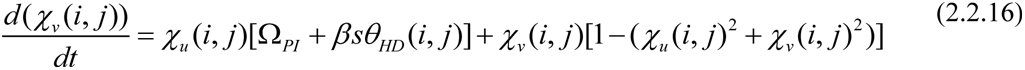

χ_u_ and χ_v_ are the u and v components of the phase of the oscillatory neuron which encodes the PI value.

Ω_PI_is the base frequency of the PIoscillatory neuron.

*β* is the spatial scaling parameter

*s* is the speed of the animal. Speed is obtained from the average of the proprioceptive inputs from the four limbs.

*s* = *mean* (p_FR_,p_FL_,p_HR_,p_HL_)

Since p_FR_,p_FL_,p_HR,_ p_HL_are the peaks of the limb oscillations (Eqn.2.2.8 – 2.2.11) which are nothing but *θ*_*m*_(1− *kd*), *θ*_*m*_(1+ *kd*), *θ*_*m*_(1− *kd*), *θ*_*m*_(1+ *kd*) respectively, the average value of these proprioceptive pulses is *θ*_*m*_which is directly proportional to the step size (Eqn. 2.2.5) which inturn gives the notion of speed of the animal. [The derivation of oscillatory dynamics from polar form to Cartesian form is shown in Appendix A4].

The outputs of PI layer are passed as inputs to a subsequent layer dubbed asLateral Anti Hebbian Network_Spatial Cell(LAHN_SC_)layer.

#### Lateral Anti-Hebbian Network (LAHN)

LAHNis a neural network with Hebbian forward connections and anti-hebbian lateral connections, known to be capable of extracting sparse features from the intput patterns(Foldiak, 1989). LAHN response is computed as follows.

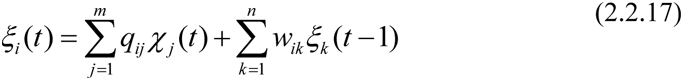

*ξ* is the output of LAHN.

*χ* is the input PI value

q_ij_ is the forward Hebbian connection from j^th^ input to i^th^ output neuron.

w_ik_ is the lateral connection between i^th^ and k^th^ output neuron.

*m* is the input dimension.

*n* is the number of output neurons.

Lateral and forward weight update equations are given as,

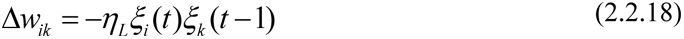

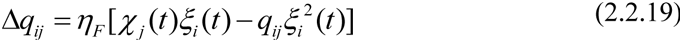

η_L_ is the learning rate for lateral weights

η_F_ is the learning rate for forward weights.

After training, weights of the LAHN_SC_ converge to the subspace of the PCs of the input vectors(Foldiak, 1989). As described in the simulations of results section(Fig 5B1-5.E.1.), neurons of LAHN_SC_ developed unique spatial firing fields resembling hexagonal and square grid cells, border cells etc.

**Fig 5:**
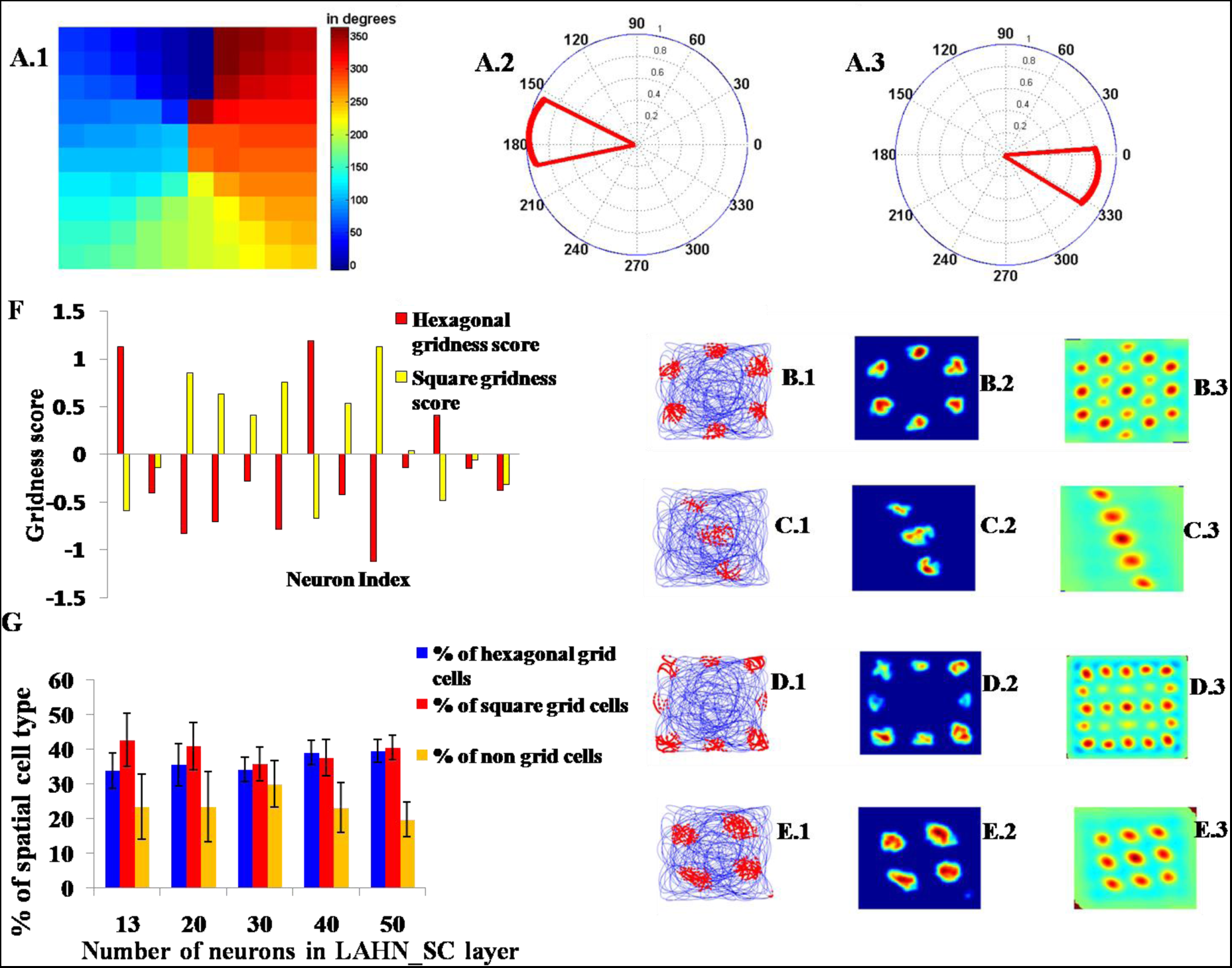
(A.1) The HD map formed from HD layer which shows topographic organization of head directions. (A.2-3) Polar plot showing the direction tuning of two neurons located at [4,1] and [7,7] in the HDlayer(A.1). (B-E) The firing field (B.1-E.1), firing map (B.2-E.2) and the autocorrelation map (B.3- E.3) of the LAHN_SC_ neurons (F). Graph showing the gridness scores (hexagonal and square) of the neurons of the LAHN_SC_ layer. Those neurons with hexagonal gridness score > 0 are regarded as hexagonal grid cells and those with square gridness score >0 are regarded as square grid cells. Those cells with both gridness scores < 0 do not fall into any of the 2 categories. (G) Graph showing the percentage of spatial cell types formed as the number of neurons in the LAHN_SC_ layer is varied.

A second LAHN named as LAHN_PC_of smaller size is placed on the top of the LAHN_SC_and LAHN_PC_ is trained using the responses from LAHN_SC_. Training equations for LAHN_PC_ are the same as that of LAHN_SC_(eqns. 2.2.18 and 2.2.19). After training, LAHN_PC_ neurons began to show localized firing field activity analogous to that of place cells (see resultssection Fig 8A.1-C.1.).

**Figure 8:**
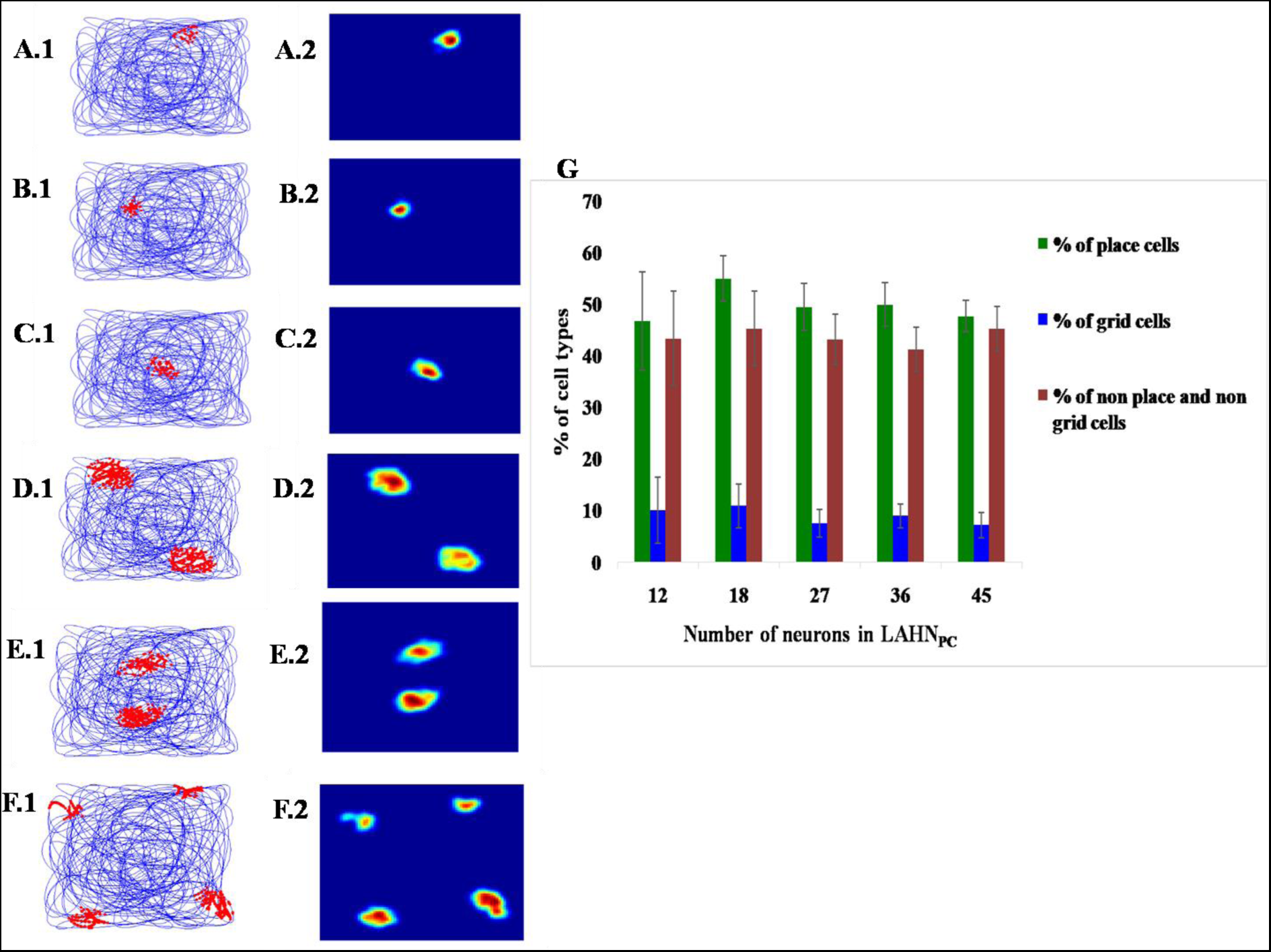
(A) Firing fields of neurons in LAHN_PC_ layer (A.1-F.1) and its corresponding firing rate map (A.2-F.2)(G) Characterization of LAHN_PC_ layer by graphing the %of cell type vs number of neurons in the layer.

### 2.3 Measure of Gridness

Both hexagonal and square gridness of the firing fields of each neuron in LAHN_SC_ are quantified using hexagonal and square gridness scores respectively. These are computed from the autocorrelation map which is obtained using equation (2.2.20) below(Hafting et al., 2005). After the computation of autocorrelation map, hexagonal and square gridness scoresare computed using equations (2.2.21) and (2.2.22).

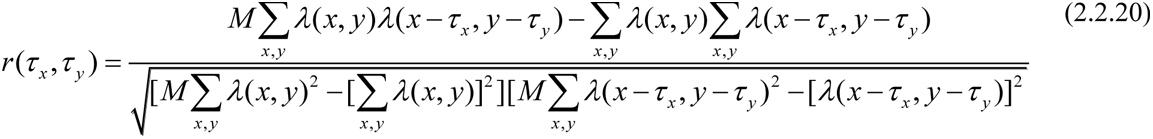

r is the autocorrelation map.

λ(x,y) is the firing rate at (x,y) location of the rate map.

M is the total number of pixels in the rate map.

τ_x_ and τ_y_ corresponds to x and y coordinate spatial lags.

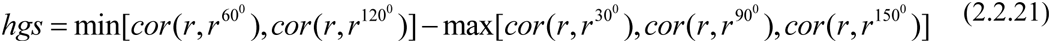

hgs is the hexagonal gridness score (HGS)

r^θ^ is the rotation of autocorrelation map by θ degrees.

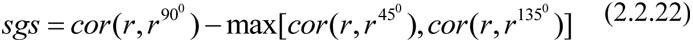

sgs is the square gridness score.

### 2.4 Phase precession

Phase precession is analyzed when the animal enters and exits a place or grid field. The phase of the spikes precesses with respect to the background theta rhythm as it traverses the field. This is quantified in terms of the regression that is done on the grid or place cell spiking phase data. If the fitted regression line has a negative slope, it indicates that the phase is precessing as the animal traverses the respective firing field(O’Keefe and Recce, 1993).

Phase precession is typically studied in 1-dimensional firing fields, which is much simpler than phase precession in 2-dimensional firing fields. We consider both forms of precession in the current modeling study. To model 1D phase precession, the virtual animal was made to move along a one dimensional track. A regression line was fitted on the phase of firing of grid and place cells from LAHN_SC_ and LAHN_PC_ respectively with respect to the constant background theta oscillation of 6 Hz whenever the animal traverses the firing field.

In order to characterize the phase precession in a 2Dfiring field, the orientation of the animal’s trajectory within the firing field must be considered. Two parameters characterize the animal’s trajectory within the firing field: the orientation, *θ*_trj_, of the trajectory (assumed linear) within the firing field, and the distance of the trajectory, *r*_*trj*_, from the center of the firing field (Fig. 2A). Fixing the orientation, *θ*_trj_, we change the distance, *r*_*trj*_, in small steps and consider phase precession. Once the animal exits the firing field it renters at a different *r*_*trj*_ but same *θ*_trj_.(Details onconstructing such a trajectoryare given in Appendix A1.2). Once it spans the desirable range of ‘r_trj_’, the virtual animal was made to reenter the firing field with different *θ*_trj_, while scanning the same range of ‘*r*_trj_’. This is repeated until the virtual animal spans all the *θ*_trj_ and *r*_trj_ which gives rise to a family of parallel trajectory segments that traverse the grid field at various *r*_trj_, but at a constant *θ*_trj_ (Fig. 2B).The difference between the spiking phase (with respect to a constant 6Hz theta rhythm),Δ*φ*_trj_, at the entry and exit of the firing field gives a measure of how much the phase precesses during its traversal at a specific *θ*_trj_ and *r*_trj_. The computed Δ*φ*_trj_ was polar plotted using *r*_trj_, *θ*_trj_ as shown in the results section.

**Fig. 2:**
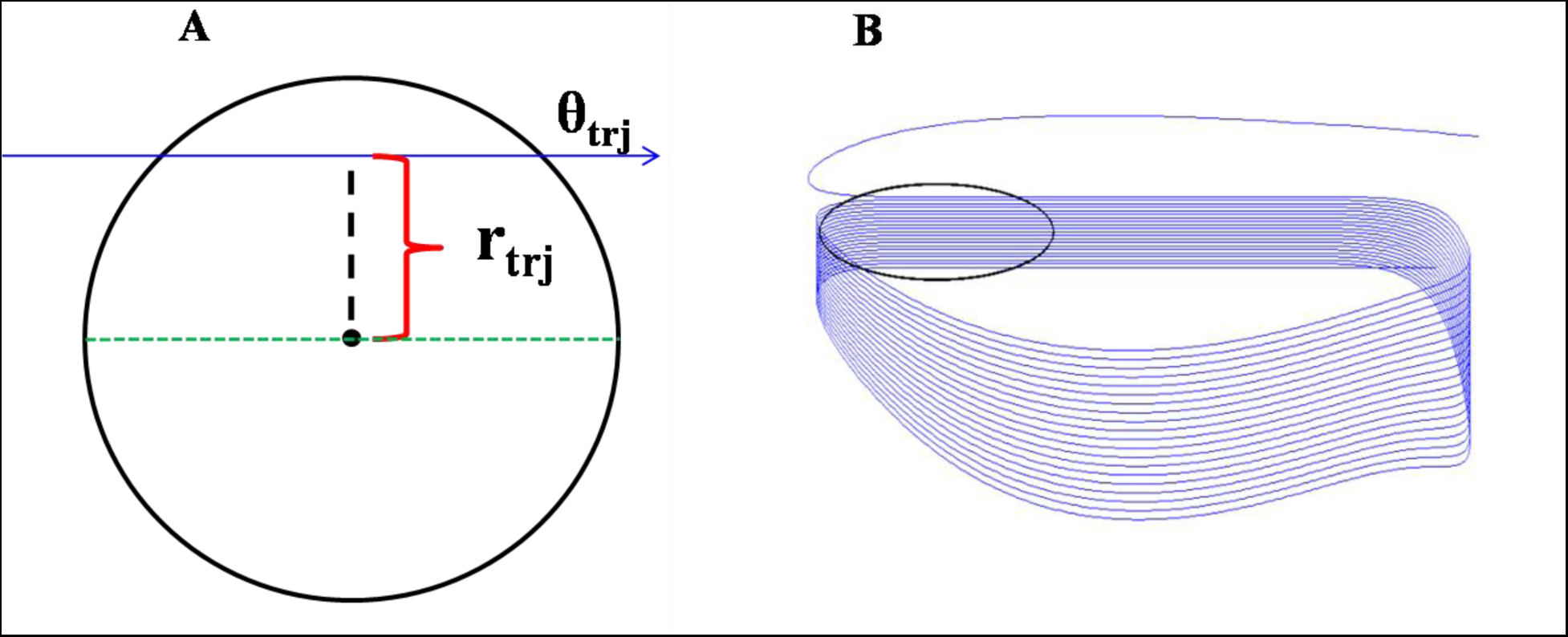
(A) Blue line indicates the trajectory traversed by the animal at a specific direction *θ*_trj_ and a distance of *r*_trj_ through a firing field (place/grid cell) indicated as black circle. Green line indicates the line parallel to the current running direction and passing through the centre of the firing field from which *r*_*trj*_ is computed. (B) A sample trajectory traversed by the animal through the firing field (black circle) that spans all the *r*_trj_ at a specific *θ*_trj_.

## Results

### A. Results from Model 1

The virtual animal was made to forage in a square box. The neuronal firing activity was represented in three different forms such as *firing field* of the neuron, *firing rate map* and the *auto correlation map* (Fig 3 A-F).Those positions on the trajectory at which the respective SC layer neuron activity as per eqn. (2.2.17) crosses a threshold (*ε*_*SC*_) are marked as red dots to get the firing field of the neuron. Firing rate map determines the activity (firing rate) of the neuron in its firing field. Firing activity varies from high (red) to no activity (blue) in the firing rate map. Autocorrelation map is computed from firing rate map using eqn.2.2.20. The main purpose of autocorrelation map is to decipher any underlying symmetry present in the firing field of the neuron. Heaxagonal and square gridness scores are computed from the autocorrelation map.Fig.3A (A.1-3) and fig. 3B (B.1-3) corresponding to the second and third PCshowed a regular firing pattern which could not be matched to any spatial cell type currentlyreported in the literature. Fig.3C (C.1-3) and fig. 3D (D.1-3)depict square-like grid cells, plotted using the fourth and fifth PC respectively. Fig. 3E (E.1-3) and fig. 3F (F.1-3) corresponding to the sixth and seventh PC respectively showed a regular hexagonal-like firing pattern. A quantitative analysis of the grid patterns was done by estimating the gridness scores as per eqn. 2.2.21 -22.

Analysis of the gridness score (Fig.3G) confirmed the presence of hexagonal and square grid cells in the SC layer. The neurons that have a hexagonal gridness score > 0 are considered as canonical grid cells(Hafting et al., 2005). Similarly, neurons with a positive square gridness scores are labelled as square grid cells.

Fig.3H shows the graph between the number of head direction cells (x-axis) and average Hexagonal Gridness score (HGS) (y axis). Since in model 1 two neurons (neurons that receive 6^th^ and 7^th^ PC) acted as grid cells, average HGS of those neurons alonewascomputed for this plot. This analysis was done to show that the hexagonal firing fields formed are robust and independent of the number of head direction cells. The model was tested under different cases where the number of head direction cells varied from 50 to 1000 and the hexagonal symmetry was quantified by the average HGS as explained above. Linear regression was done on the data (p<0.001) and the obtained slope was very small (m=0.000089) which shows that the variation in the number of head direction cells does affect the hexagonal gridness scores of the neurons significantly.

### B. Results from Model 2

The proprioceptive pulses (eqn. 2.2.8-11) which are produced using the curvature-modulated locomotor rhythms (eqns. 2.2.1-4) were integrated by the oscillatory network (eqn. 2.2.12-13), whose outputs are passed on to the HD layerto obtain the HD map. Fig. 4 shows a short curved segment of the trajectory (Fig. 4A),the corresponding curvature-modulated limb oscillations (Fig. 4B) and proprioceptive pulses (Fig. 4C) respectively.

**Fig. 4:**
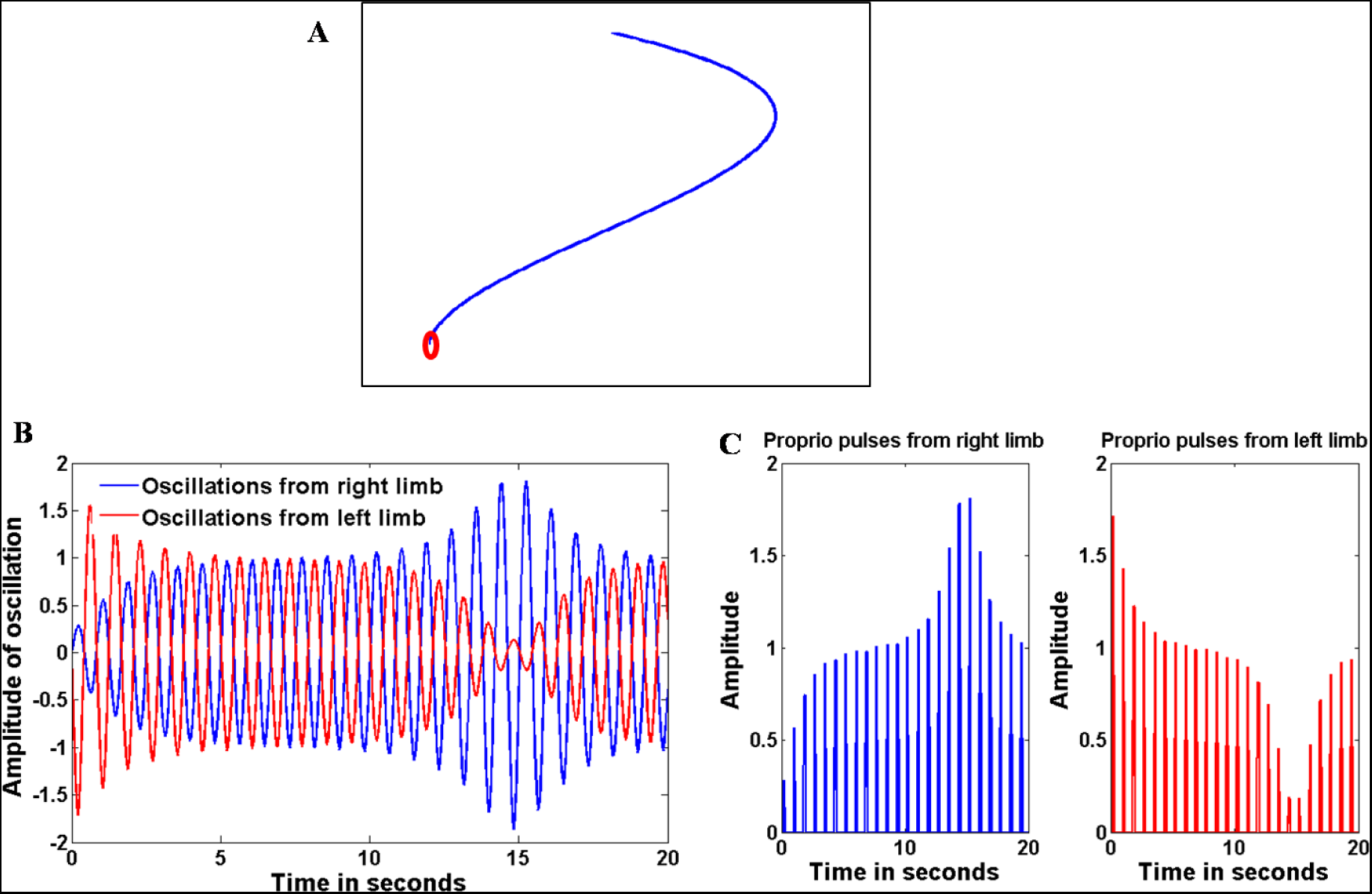
(A) A simple curved trajectory (red circle indicates the starting position), (B) the corresponding limb oscillations whose amplitudes are modulated by the curvature of the trajectory, and (C) the proprioceptive pulses extracted from the limb oscillations. In order to characterize the head direction tuning of HDneurons, the activity of that respective neuron (eqn. 2.2.14), whenever it crossed a threshold (*ε*_*HD*_), was polar plotted as a function of the current direction of motion (Fig. 5A.2-3). Fig. 5A.2-3 shows the direction tuning of the HD neurons. Fig 5A.1 shows the image plot of the weights of the HD neurons which shows a topographic organization of directions on the HDlayer.

Beyond the HD layer another layer of oscillatory neurons, the PI layer, which is of the same size (10x10) of the HD layer, performs path integration. Phases of these PI oscillators were integrated based on the current head direction of the animal (eqns.2.2.15-16), represented as a unique activation pattern on the HD layer(eqn. 2.2.14).

Training LAHN_SC_ network on the output of PI oscillators gave rise to cells with spatially periodic firing fields (Fig 5B-E). As in the Model 1, Model 2 also gave rise to varioustypes of spatial cells but the heterogeneity in the observed firing fields was greater in the case of Model 2. Apart from hexagonal andsquare firing fields, border-like activity (Fig 5D.1) was also observed in Model2.

Like in Model1, we calculated the gridness scores for all the neurons in LAHN_SC_ (fig. 5F). Percentage of cell types formed using different number of neurons in LAHN_SC_ layer was verified as shown Fig 5G. This graph was obtained by retraining the LAHN_SC_ network 20 times using N number of neurons. For N number of neurons in LAHN_SC_ layer, an average of ≈ 37% of neurons formed hexagonal grid cells, 39.6% formed square grid cells and ≈ 24% formed non-grid cells that include border cells and cells with random firing fields. This concurs with the experimental observation (Couey et al., 2013; Hafting et al., 2005; Krupic et al., 2015; Solstad et al., 2008) that EC has all the aforementioned spatial cell types with grid cells being the major spatial cell (≈50%) and other cells like border cells to be fewer in number (≈ 10–15%)(Solstad et al., 2008). In our model we gave the statistics of separate grid cells (hexagonal and square) and non-grid cells (inclusive of border and random firing cells). We did not quantify border cells separately like how we did for grid cells. Irrespective of the number of neurons in the LAHN_SC_we observed similar kind of heterogeneity in these different populations of neurons.

#### Spatial characteristics of grid cells

We also analyzed the gradient in the grid scale seen along the dorso-ventral axis of the medial entorhinal cortex (mEC). Grid scale is the distance between the central peak and one of the vertices of the inner hexagons of the autocorrelation map (Brun et al., 2008; Stensola et al., 2012). Studies have shown that (Brun et al., 2008; Stensola et al., 2012) the grid scale increases from dorsal to ventral mEC. In order to capture this trend we varied the ‘spatial scale parameter,’ β, in the PI equation (eqns. 2.2.15-16). β determines the intrinsic oscillation frequency of the path integration oscillatory neuron. For two neurons with different β such as β_1_> β_2_, that neuron with β_1_ oscillates at a higher frequency than that with β_2_ for the same speed (*s*) and head direction response (θ_HD_). This makes the first neuron to reset faster than the second and hence this gets reflected in the spatial scale of the grid cells. Variation of grid scale as a function of β is depicted in Fig 6A. Grid scale is quantified in terms of minimum and median grid scale (Brun et al., 2008; Stensola et al., 2012). In both cases, distances of the six inner hexagon vertices from the central peak of the autocorrelation map are computed. The minimum value among of these distances represents the minimum grid scale and the median value represents the median grid scale. This quantification is shown in Fig 6B.1 and also compared with the experimental results (Fig 6B.2).

**Figure 6:**
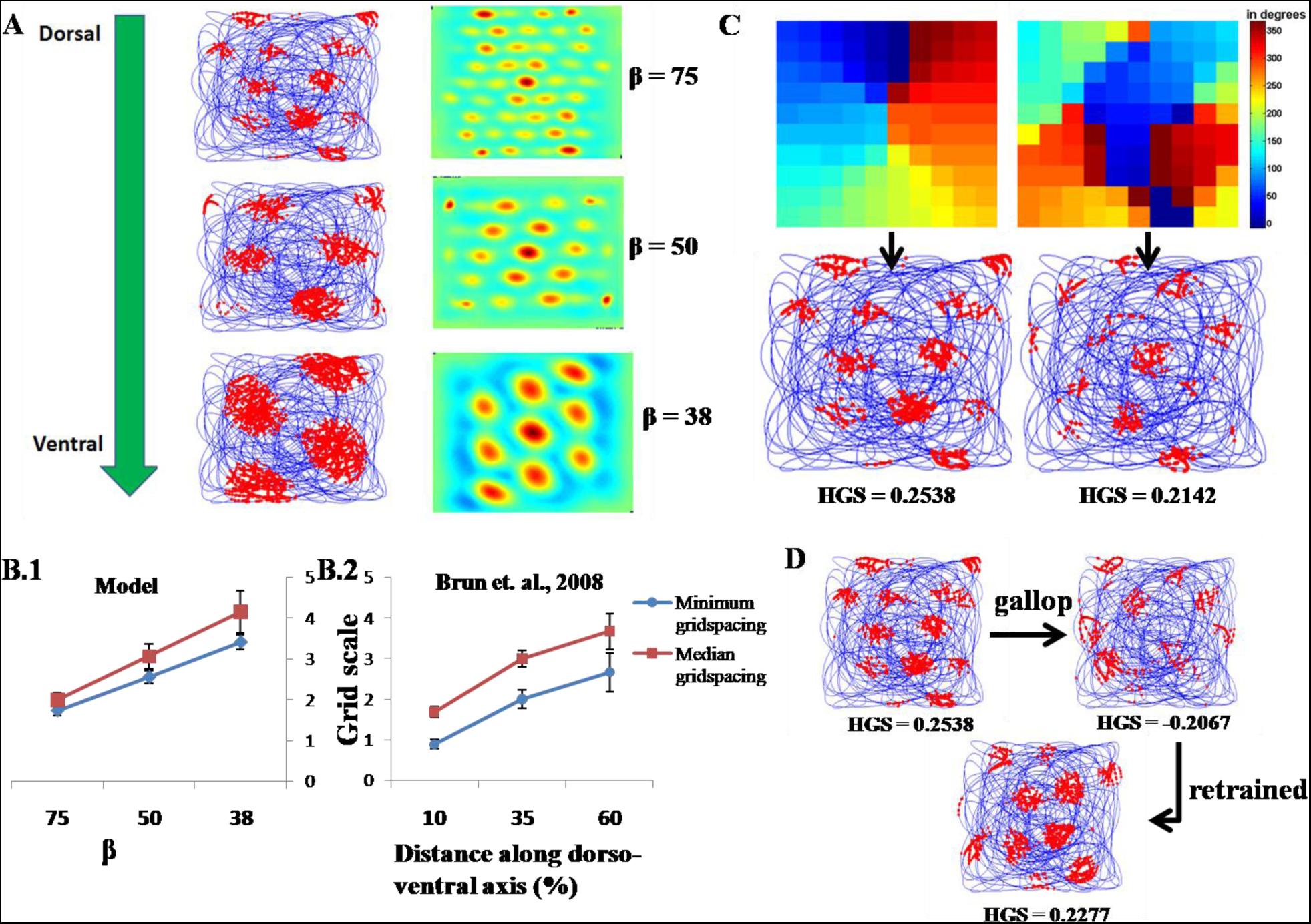
(A) Dorso-ventral gradient of the grid scale is captured by the model by changing the spatial scale parameter β.(B) Comparison between the model (B.1) and the experimental (B.2) quantification of grid scale along the dorso-ventral axis of mEC.(C) Formation of grid fields with and without topographic organization on the HD map. (D)The change in the grid firing field on changing the locomotor rhythm from trot to gallop.

There are instances in literature where the existence of topographic maps in the HD cell layer has been contested(Giocomo et al., 2014; Taube et al., 1990a). Inorder to examine the effects of topographic arrangement of HD cells and the resulting formation of the grid cells, we varied the training parameters of the HDlayer algorithm (the radius of neighborhood) to produce maps with different characteristics. Fig 6C shows both a smooth topographic map of the HD cell layer (obtained by using a neighborhood of 3) and the HD layer without a distinct map structure (obtained with smaller neighbourhood of 1).In both the cases, training the LAHN_SC_ layer gave rise to hexagonal grid cells, suggesting that topography in HD layer may not be a determining factor for grid cell formation. Additionally studies in rats show that there is no topography seen in post-subiculum as the electrode is moved obliquely through the postsubiculum (Taube et al., 1990a). The corresponding grid fields for both the HD maps are indicated by the arrows (Fig 6C).

We also studied the effect of changing the locomotor rhythms and their influence on the evolution of grid fields. We considered two types of rhythms which are normally seen in animals when shifting their movement from walking (trot) to running (gallop). This change in rhythm was brought about by changing the phases of the limb oscillators(Collins and Richmond, 1994). Whereas in trot the contralateral limbs are in phase, in gallop both the forelimbs are in phase and out of phase with the hind legs. On changing the rhythms from trot to gallop, we saw that there was a loss of grid field formation (Fig 6D). But on retraining the network on the new rhythm, grid fields reemerge. This could suggest that on changing their rhythms, animals might have to remap their spatial representations in order to account for the changes in patterns of sensory (in this case the proprioception) inputs coming to the EC. Another possibility is that, perhaps, separate sets of grid cells respond to distinct locomotor rhythms.

#### Temporal characteristics of grid cell

The temporal code underlying grid cell activity has been explained in terms of phase precession, which refers to the gradual shift in the phase of firing of a grid cell with respect to the background theta rhythm as the animal progresses along a track (Hafting et al., 2008). There is a characteristic variation in the phase as the animal passes through a grid field, which is described as a marker by which the animal can estimate its position inside the field. The model was tested to understand whether the same phenomenon of phase precession emerged after training the network. Phase precession studies were done only with Model 2 and not with Model 1 because of the greaterbiological plausibility of Model 2.

In experimental and modeling literature, phase precession is often described when the animal moves along a 1D track (Blair et al., 2008; O’Keefe and Recce, 1993). However, since grid fields are 2D, it is essential to characterize phase precession with respect to 2D grid fields. There are instances in the literature where 2D phase precession is analyzed using circular regression (Jeewajee et al., 2014). Here we follow a different method. In the present study, we present a method of characterizing phase precession in 2D grid fields as described in the methods section 2.4.

First we describe phase precession results as the virtual animal moves along a 1D track. Fig 7B shows the phase precession of grid cell on the 1d track and is evident that phase of the grid cell precesses with respect to the theta rhythm.

**Fig 7:**
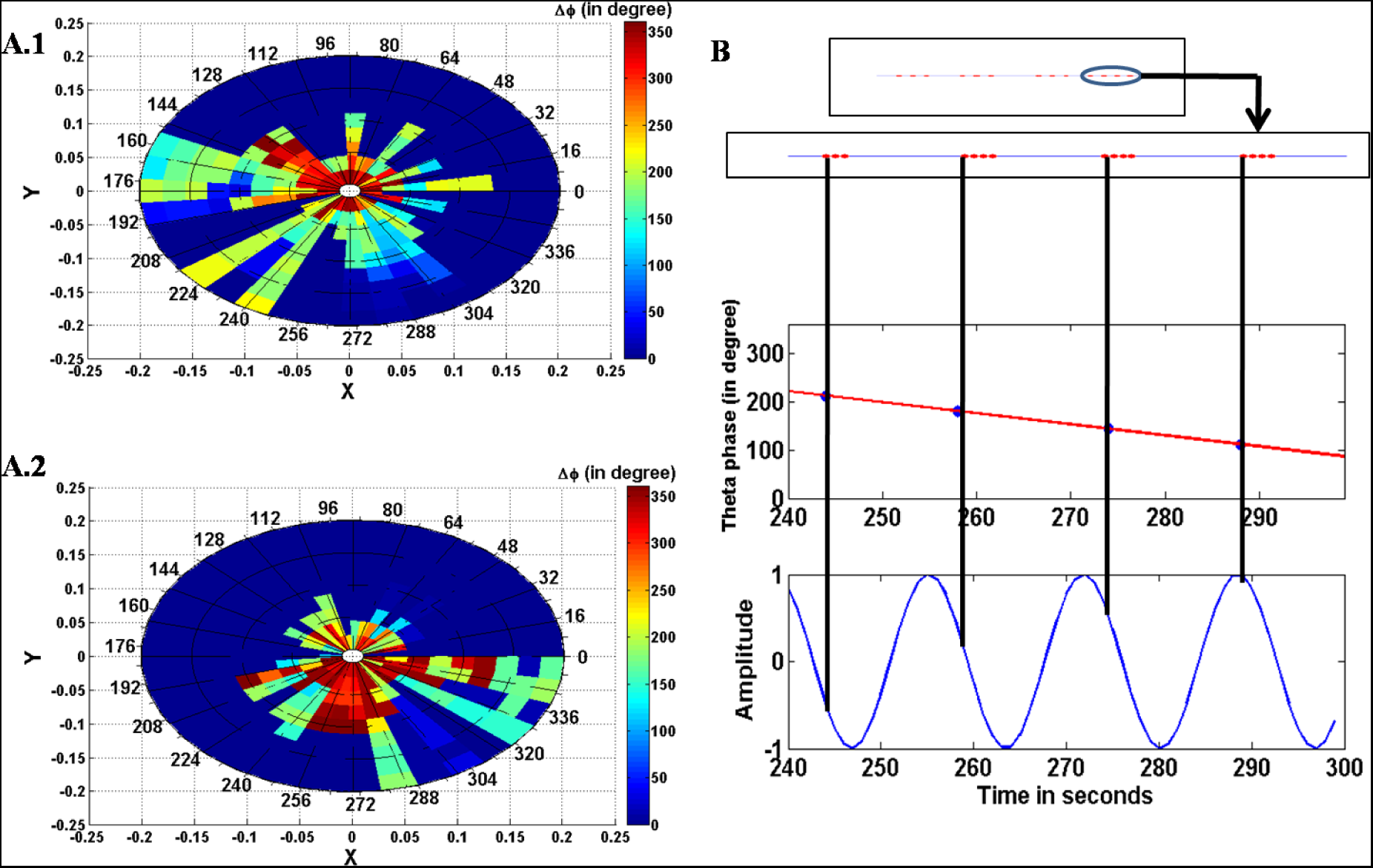
(A.1) Polar plot of Δ*φ*_trj_ as a function of X [*r*_*trj*_*cos(θ*_*trj*_)] and Y [*r*_*trj*_*sin(θ*_*trj*_)] for clockwise *θ*_*trj*_ through the grid field.(A.2). Polar plot of Δ*φ*_trj_ as a function of X [*r*_*trj*_*cos(θ*_*trj*_)] and Y [*r*_*trj*_*sin(θ*_*trj*_)] for anticlockwise *θ*_*trj*_ through the grid field.(B) Phase precession of grid cell while the animal moves on a 1d track. Blue oscillations represent the constant background theta wave. Linear reduction in phase as the animal moves forward can be noted.

In order toquantify phase precession in 2D firing field, we express Δφ_trj_ as a function of θ_trj_ and r_trj_. Fig 7A.1 and A.2 shows the polar plot of Δφ_trj_ as a function of X [r_trj_cos(θ_trj_)] and Y [r_trj_sin(θ_trj_)]. Since the grid field radius was 0.2, in the polar plot, *r*_trs_ ranges from -0.2 to 0.2 as shown in Fig 7A.1 and 7A.2. Note that when Δφ_trj_is depicted as a function of r_trj_andθ_trj_, for a given r_trj_andθ_trj_, the virtual animal can move in clockwise or anticlockwise direction.Fig 7A.1 shows the polar plot for θ_trj_, with the virtual animal moving in clockwise direction and Fig 7A.2 shows the polar plot for θ_trj_, with the virtual animal moving in anticlockwise direction. From fig.7A.1 and fig.A.2 it is evident that Δφ_trj_ is maximum when r_trj_ is small i.e. when the animal is close to the centre of the grid field. However, variation of Δφ_trj_ with respect to r_trj_andθ_trj_ is anisotropic because its variation is not the same in all directions, θ_trj_, which is evident from Fig 7A.1 and A.2.This means that the grid field firing is direction-dependent; otherwise both plots must be identical. Hence it suggests that grid field activity could encode, while the animal is inside the grid field, not only the location of the animal but, perhaps, also its average direction of motion within the grid field.

#### Emergence of place activity

The model was extended to understand whether another LAHN layer could give rise to cells thatare higher above in the hierarchy in the HC. The output of the LAHN_SC_ layer (eqn. 2.2.17) was passed onto a second LAHN_PC_ layer (LAHN_PC_). Like LAHN_SC_, training of LAHN_PC_ is also done following eqns. (Eqn. 2.2.18 and 2.2.19). Remapping the response of the neurons in the LAHN_PC_ layer on the trajectory showed a highly localized spatial firing field. This response resembles that of the place cell that fires only at a specific location known as place field in the two dimensional space. Fig 8A shows the place cell-like 2.2.19 response of the neurons in the LAHN_PC_ layer. Anatomical studies report that(Akdogan et al., 2002; Mulders et al., 1997) the ratio of number of neurons in hippocampus CA1 to EC (considering only layer 2 and 3 of MEC since they form major synapses with CA1 (Moser et al., 2014)) in rats is ≈ 0.91. Hence in the model also we followed the same ratio of number of neurons of LAHN_PC_ to LAHN_SC_ layer. The pertinent feature that we noticed in LAHN_PC_layer after trainingis that neurons in this layer showeda tendency tohave localized firing fields. For characterizing LAHN_PC_ layer we followed the same method that we adopted for LAHN_SC_ layer i.e. by changing the number of neurons in LAHN_PC_ (proportional toLAHN_SC_) and measuring the composition of various cell types that evolved. To characterize the place cell we examinedthe number of peaks in the autocorrelation map (Eqn. 2.2.20) of the respective LAHN_PC_ cell. If the number of peaks is one, then the cell is characterized as a place cell. This is because, a place cell has localized firing field and lacks spatial periodicity. Hence its autocorrelation map will have only one single peak unlike spatially periodic grid cells which have multiple peaks in its auto correlation map. Fig 8.A.1 – F.1 shows the firing fields of the cell types found in LAHN_PC_layer. Fig 8A.2.-F.2 shows the respective firing rate maps. Fig 8.G shows the characterization of LAHN_PC_ layer. From the graph (Fig. 8.G) it is evident that an average of 49.73% of LAHN_PC_ neurons evolve as place cells, 8.85% evolve as grid cells (hexagonal or square) and 43.65% evolve as non-place and non-grid cells but showed firing fields with less periodicity like corner cells. These results suggestthat as the hierarchy of the unsupervised layer increases the neurons show more “placeness” than “gridness”.

Additionally the place cell’s temporal characteristics were ascertained using a strategy similar to the one used for the grid cells. Fig.9A shows the phase precession of a place cell from the LAHN_PC_ layer on a 1D track. Phase precession is evident from the negative slope of the regression line.Fig.9B.1 and fig.B.2 show the polar plot of Δ*φ*_trj_ as a function of X [*r*_*trj*_*cos(θ*_*trj*_)] and Y [*r*_*trj*_*sin(θ*_t*rj*_)] for clockwise and anticlockwise θ_trj_. In the case of place cell also Δ*φ*_trj_ is maximal close to the centre of the grid field for both clockwise and anticlockwise θ_trj_. Fig 9B.1 and fig.B.2 also show anisotropy in place cell coding.

**Figure 9:**
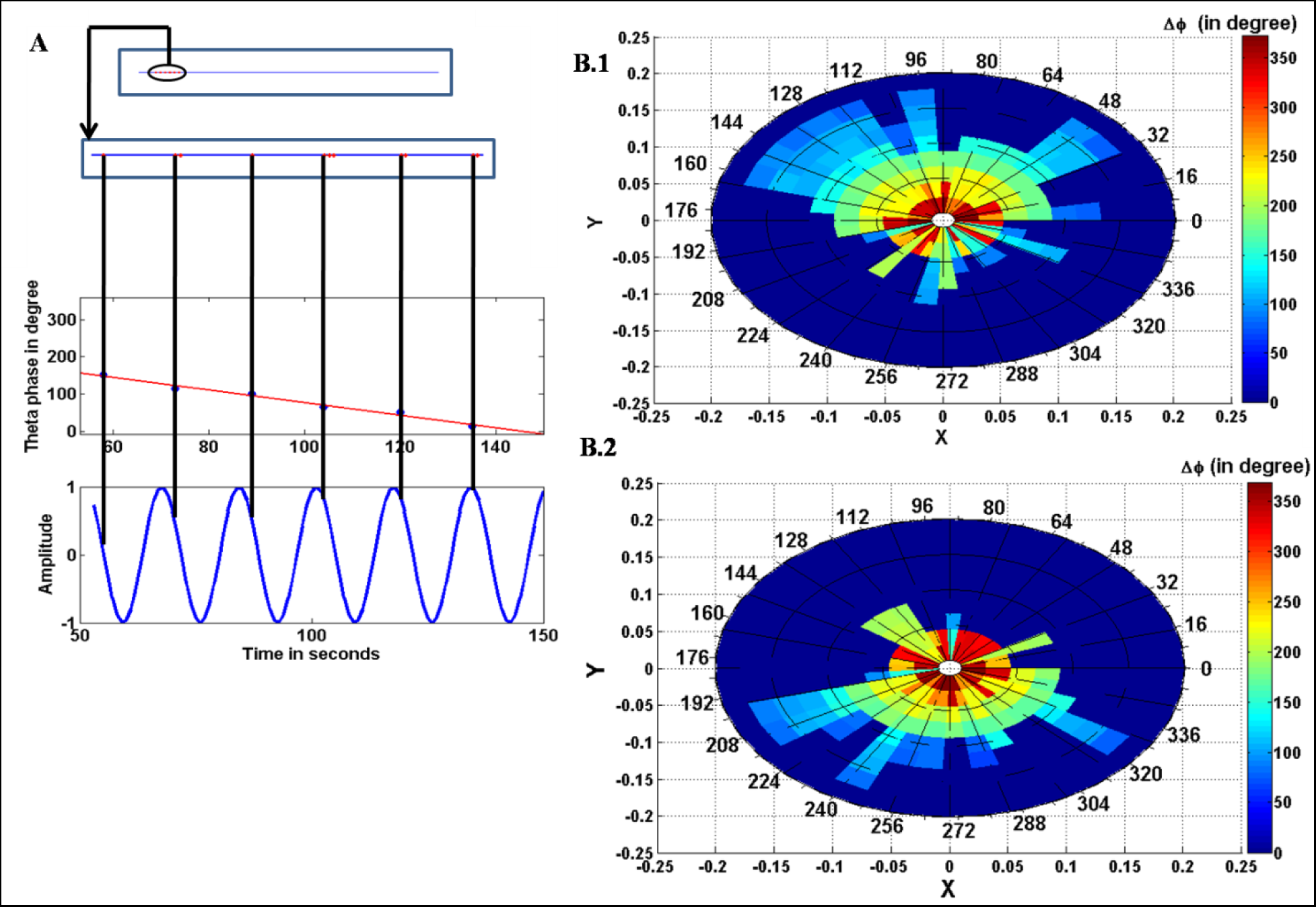
(A) Phase precession of place cell tested while the animal was made to navigate on a 1D track. (B.1) Polar plot of Δ*φ*_trj_ as a function of X [*r*_*trj*_*cos(θ*_*trj*_)] and Y [*r*_*trj*_*sin(θ*_*trj*_)] for clockwise *θ*_*trj*_ through the place field. (B.2)Polar plot of Δ*φ*_trj_ as a function of X [*r*_*trj*_*cos(θ*_*trj*_)] and Y [*r*_*trj*_*sin(θ*_*trj*_)] for anticlockwise *θ*_*trj*_ through the place field.

## Discussion

We proposed a biologically plausible model for the spatially tuned hippocampal cells like head direction, grid (both hexagonal and square), border and place cells in a hierarchical fashion using locomotor rhythms as the input. One of the key mysteries of grid cells is the origins of hexagonal symmetry, when such symmetry exists neither in the inputs nor in the networkarchitecture itself. We started with Model 1 in which it was possible to show that the PCs associated with the PI layer have special periodicities easily related to grid cell periodicities – square and hexagonal. PCs in which the peak-to-peak angular deviation is 60 degrees correspond to hexagonal grid cells; those in which the peak-to-peak angular deviation is 90 degrees correspond to square grid cells (Figures shown in Appendix 2).

Existing models of grid cells often make special symmetry assumptions which tend to weaken the biological significance of the model. One such example is the assumption of existence of head direction cells whose phases are separated conveniently by integral multiples of 60^°^ which is biologically rathercontrived(Burgess et al., 2007; Hasselmo et al., 2007;Zilli and Hasselmo, 2010). Our Model 1 was able to circumvent this, by showing that grid cell symmetries have their roots in the symmetries of the eigenvectors associated with theresponses of the PI layer. The robustness of the emergence of hexagonal grid cells from the proposed model is established by the fact that the HGS score is quite robust against variation inthe number of HD cells (Fig 3H).

The proposed model has another advantage over several existing grid cell models. Unlike existing models that are often specially designed to exclusively exhibit hexagonal grid cell responses, our model naturally exhibits grid cells of several kinds of symmetries – hexagonal and square and also cells that fired at the corners of the space. Furthermore, these grid cells do not have perfectly hexagonal periodicities which would also be biologically unrealistic: their hexagonal (or square) gridness score varies over a considerable range (Fig 3H).

Another interesting feature of our modeling approach is the manner in which HD cells are modeled. HD layer is often modeled as a ring of cells forming a 1D continuous attractor (Gaussier et al., 2007; Knierim et al., 1995; McNaughton et al., 1991; Redish et al., 1996; Sharp et al., 2001; Stringer et al., 2002; Zhang, 1996). Existence of special rings of neurons in the brain is a difficult assumption to confirm. Instead in our model, HDcells form a 2D layer of neurons. Furthermore, this 2D layer of HD neurons may or may not have map structure, a feature that has no bearing on evolution of spatial cell responses downstream.

Previous models implemented integration of the so called angular head velocity (AHV) signals in a biologically unfeasible way. For instance Zhang*et al*(Zhang, 1996)assumed symmetrical and asymmetrical connections amongthe head direction cells where the asymmetrical connections were modulated by the AHV signals. These fast changes in synaptic connections precisely modulated by AHV signals are biologically unfeasible. Also some models (Goodridge and Touretzky, 2000; Redish et al., 1996; Stringer et al., 2002;Zhang, 1996)consist ofnetworks of HD cells,hard-wired along with the so-called ‘rotation cells.’ These rotation cells receive the AHV signals and move the stable activity packet along the ring of HD cells. In the present model,the direction of motion is computed by measuring the difference in the limb oscillations on the ‘inner’ side as opposed to the ‘outer’ side as the virtual animal makesa turn; this differenceis further integrated into the phase of respective oscillators. In the proposed model, path integration is achieved by integrating the head direction inputs and speed into the phase of an internal oscillation.

An essential result in the proposed modeling approach is that principal components extracted from PI neural responses lead to cells with grid-like periodicities (square, hexagonal etc.). It may be commented that principal component analysis is a slightly artificial data analysis method without a strongneural basis. Although there are neural architectures that can extract top K principal components, these are of a rather artificial construction (Oja, 1982; Sanger, 1989). Therefore, we replaced the PCA stage in Model 1, with the LAHN stage in Model 2. LAHN requires only lateral anti-hebbian connections and afferent hebbian connections in the output layer (Foldiak, 1989). Both these forms of synaptic plasticity are biologically feasible and are found in many different parts of the brain (Couey et al., 2013).This data driven neural network with n number of output neurons converges its weights to the subspace of the first n principal components as its basis vectors.

Grid cells, which are typically found in EC, are one synapse upstream of place cells which are predominantly seen in the CA1 region of the hippocampus (Moser et al., 2014).Following the same hierarchy,we trained LAHN_PC_ aboveLAHN_SC_ such that the number of neurons in LAHN_PC_is slightly less than that in LAHN_SC_. The neurons in LAHN_PC_beganto develop localized firing fields akin to place cells after training. This seemed to indicate that all the spatial periodic cells which have different firing fields are necessary to initiate the evolution of a place cell. This possibly makes sense as the animal would require complete information about its space which we presume is available in the LAHN_SC_ layer, before mapping them onto specific salient locations in the environment. Formation of place-like activity happened naturally without any constraints in the model. Hence this model was able to explain the possible mechanism of formation of place cells from proprioceptive information. Place cells are well anchored to distal cueslike visual landmarks(O’keefe and Nadel, 1978), in such a way that any manipulation to the stationary landmarks like rotation or translation gets reflected in the place field map also. Since we did not incorporate any visual inputs to the model, we could not analyze the place cells in terms of their visual responses.

Experimental literature (Brun et al., 2008; Stensola et al., 2012) showed that there exists a gradient of grid scale along the dorso-ventral axis of the entorhinal cortex such that it varied from low to high value from dorsal to ventral region respectively. In order to take this into account in the model, we varied the spatial scale parameter β in path integration (Eqns.(2.2.15-2.2.16). βdetermined the intrinsic oscillation frequency of the path integration oscillatory neuron. Giocomo et al (Giocomo et al., 2007) had already shown empirically that the sub-threshold membrane potentialoscillation frequency decreases from the dorsal to ventral region of the 2^nd^ layer of the medial entorhinal cortex. Hence those dorsal neurons with higher intrinsic frequency get quickly reset during path integration and can result in smaller spatial scale, with the opposite trend seenin case of ventral neurons. Burgess and group in theiroscillatory interference model (Burgess et al., 2007)also used the same notion for the spatial scale parameter.

Taube *et al* (Taube et al., 1990a) showed that there was no trend for the neighboring HD cells to have similar preferred head direction. This means thatthe HD layer lacked topography. We were able to produce head direction models without topography by retraining the HD layer with a smaller neighbourhood radius. Even though the fields get remapped, loss of topography in the head direction map does not affect the grid formation in the LAHN_SC_ respectively as shown in Fig. 6C in the results section.

Hence the proposed model gives more insight into the possible mechanism of spatial cell formation even in the absence of topography in the head direction maps, as shown empirically.

Experimental studies show that grid cells are more dependent on the proprioceptive inputs since they retain their hexagonal firing field even after the removal of stationary cues like visual landmarks or olfactory cues (Moser et al., 2014). Therefore, we were interested in analyzing the effect of proprioceptive locomotor rhythm on the grid field formation. To our knowledge, this is the first attempt, in computational terms, to analyze the impact of locomotor rhythm on the formation of grid cell responses. Two rhythms such as trot and gallop were considered as examples of locomotor rhythms. We did not model a comprehensive locomotor gait pattern generation; instead we changed the phases between the limb oscillators for generating respective gait like rhythms. The virtual animal was assumed to have four limbs. The trot-like rhythm was simulated by making the contralateral limbs to be in phase and out of phase with respect to the other two limbs. The gallop-like rhythm was simulated by making the two front/hind limbsto be in phase and out of phase to the other limbs. Initially the model was trained using proprioceptive inputs coming from the trot-like rhythm. Once the grid fields were obtained from the respective neurons in the LAHN_SC_, the model was presented with proprioceptive inputs from gallop like rhythm. Now the previously formed grid fields were weakened or lost as shown in Fig6D. However, when themodel was retrained using the new gallop-like rhythm, the grid fields re-emerged but had a different hexagonal firing field. This remapping suggests that there are perhaps different grid cells responding to different locomotor rhythms, a strong prediction that emerges from the current model.

With special attention to grid and place cells, we analyzed the temporal characteristics i.e. phase precession of both types of neurons in one dimensional and two dimensional firing fields. Our attempt was to build a comprehensive model that brings all the spatial neurons pertaining to spatial navigation of an animal under a single roof using a biologically plausible hierarchical architecture.

We do not address the issue of causality between grid and place cells: do grid cells drive formation of place cells or is it the other way around? Rolls *et al* proved that place cells can be formed from the grid cells using competitive learning (Rolls et al., 2006). Bjerknes *et al* reported that place cells and border cells form matured firing field even at post-natal 16^th^ (P16) day when the rat pup leaves its nest (Bjerknes et al., 2014). But the grid cells were not matured during that period. This result is consistent with our modeling approach according to which locomotor inputs are necessary for grid cell formation. Place responses can be shaped by visual and other sensory stimuli also, though visual stimulus is not present in the proposed model. Furthermore, in the proposed model, place cells were formed not exclusively from grid cells; instead all the spatially periodic cells in LAHN_SC_ layer collectively contributed to the formation of place cells.

Further investigation on border cells, like manipulating the shape of the foraging environment was not done since we were mainly interested in understandingthe possible mechanism for the formation of different spatial cells. We would like to makea special note on the receptive field of the border cell from the model. The border cells in the model fields did not show any affinity towards a particular wall but fired whenever they encountered the border of the environment. Empirical results showed border cells that fired at more than one border of the environment (Stensola et al., 2012) suggesting that suchnon-specific border cells could exist in the entorhinal cortex. It is possible that visual or other sensory modalities bring this specificity to border cell responses. However, such sensory modalities are omitted from the current model. Predictions from the model can be listed as given below:

1. Our model predicts the possibility for more spatial representations other than grid or border cells like, for example, corner cells (Fig. 8D.1) that evolved from the model and encoding the corners of the environment. This is more specific than border cells which fires when the animal encounters a border. (The disappearance of corner cells needs to be ascertained in a circular pool, which will be considered as the future work)
2. Our model hypothesizes a relation between locomotor rhythms and grid cell formation (Fig 6D) and predicts the possibility of different grid cells coding for different rhythms of locomotion. General convention of using a direct velocity vector to perform path integration has been circumvented in this study by understanding the role of proprioceptive inputs obtained from locomotor rhythms to drive the formation of spatial cells. It is therefore necessary to also consider the effect of changing locomotor patterns on their representation in sensory cortices. Indeed such an effect is seen in parietal area where there is significant change in the activity of neurons in parietal area 5 to change in rhythms (Beloozerova and Sirota, 2003).
3. Our model predicts that place and grid like activity is not restricted to hippocampus and entorhinal cortex. It can even occur at higher centers of the brain like posterior parietal cortex (PPC) which the literature describes as a crucial cortical area for spatial representation and also the site of visuo-motor integration (Beloozerova and Sirota, 2003; Parron and Save, 2004; Shrager et al., 2008; Whitlock et al., 2008).

## Conclusion

We proposed a biologically plausible model of hippocampal spatial cells. The model explains the generation of head direction, spatially periodic hexagonal and square grid, border and place cells respectively from locomotor inputs. The future work includes the incorporation of visual input to the model and study the role of visual stimuli in the formation of the aforementioned spatial cells. We also like to extend our model to a full spatial navigation context wherein an animal searches for a goal location under various sensory conditions. It would be an interesting modeling exercise to study the contribution of different types of spatial cells in enabling the animal to reach its navigational goals.

## Appendix

### A1.1 General trajectorycalculation

The movement of the animalinside the square box is designed using the dynamics of curvature-constrained motion. We used a dynamical system which is used to model Dubins and Reeds-Shepp’s cars(Takei and Tsai, 2013). The dynamics are given as follows

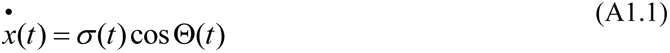

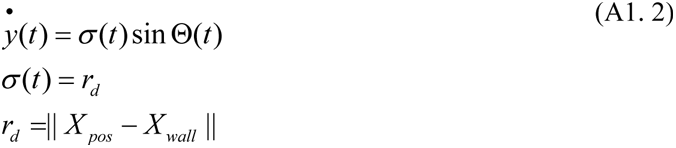

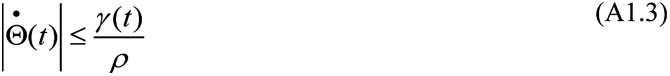

where *x* and *y* are the 2d coordinates of the animal, Θ is the direction of movement of the animal at a given time, *ρ*>0 represents the minimum turning radius of the animal and σ controls the speed of the animal. X_pos_ and X_wall_ are the Cartesian coordinates of the position and the wall at which the animal is directing. r_d_ is the distance from the animal’s current position to the nearest wallit encounters if it travelled continuously in that same direction. To ensure that the animal does not cross the boundary the speed *σ* is set as r_d_ and the animal’s speed considerably slows down as it nears the wall which is a reasonable assumption to consider for further simulations. Furthermore, in order to introduce the element of randomness in the animal’s trajectory a dynamic variable γ(t) is introduced in eqn. (A1.3). γ(t) ϵ [-1 1] is designed in such a way that there is high degree of randomness in the animal’s trajectory when the current position is far away from the walls compared to when it is close to it. A softmax rule is imposed on the probability of the animal to take a right or a left turn at a given instant. The parameter known as temperature in the softmax (here referred as *α*) is controlled by the distance of the animal to the walls. *α* is inversely proportional to the distance from the walls and its value becomes smaller and hence more random in picking the direction to move compared to when the distance to the walls is smaller and it sticks to a single direction. The dynamics of γ(t) is given by

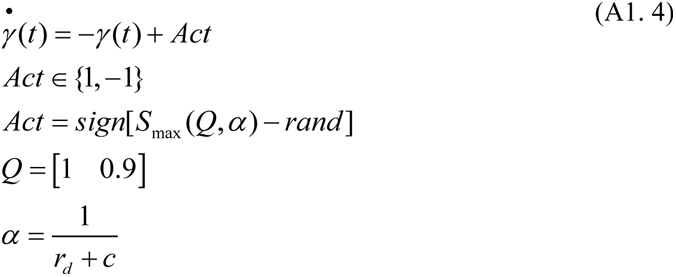

Where Act is the action to be taken, Q is the value for each action, *S*_max_ is the Gibbs softmax function which gives the probability of selecting an action which here is the direction of motion and is given as

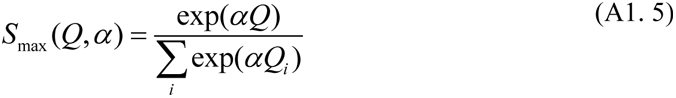

p_m_ is a parameter that switches discreetly between 1 and -1 after certain number of iterations so that the animal can turn both directions closer to the walls.

### A1.2 Special trajectory implementation

By special trajectory we meant the trajectory for analyzing 2D phase precession as shown in Fig 2. We start the special trajectory as a continuation of the main trajectory (explained in A1.1). This is to ensure that the path integration does not get interrupted in between. First we made a circle of radius *r* around the grid or place field such that the circle enclosed the single firing field of the respective cell. Hence *r* corresponds to the radius of the firing field. Next we selected fixed number of points (arranged as an array) equally spaced along a line to either side of the firing field circle. Hence each point selected at one side of the circle has corresponding partner at the other side of the circle. The animal is allowed to start from the first point at one side of the circle and allowed to traverse the firing field at a direction θ_trj_ until it reaches its partner at the other side of the circle. From there it takes a smooth turn and reaches the second starting point and this continues until it spans the entire firing field (or entire r_trj_ as explained in methods 2.2). To make the turning smooth we used *spline* function in MATLAB. The turn has to be smooth since there is an upper constraint on the curvature as explained in the methods section 2.0 (Refer Appendix A3 for more information on curvature constraint). Once this is done the array of points (at both sides of the circle) are shifted and rotated to make the animal to traverse through the firing field at different θ_trj_.

## A2. Principal Components (PC) of PI layer

PCs of PI layer (except the first PC which is a constant vector. Hence it is not taken into account) show sinusoidal nature as shown in Fig A2.A. The spatial periodicity shown by the SC layer neurons can be attributed to the peak to peak angular deviation of the sinusoidal PC weight vector that they receive from the PI layer. That PC weight vector with θ^°^ angular deviation (Fig A2.B) produces the firing field with a rotational symmetry of θ^°^ (for example if θ is 45^°^ the firing field resembles an octagonal shape. Octagon has a rotational symmetry of 45^°^). Frequency of the PC sinusoids increases as the index of the PC increases. But pair wise PCs (for example 2 and 3) show same frequency with a 90^°^ phase shift as shown in Fig A2.C. This forms the reason why the pair of SC neurons show same firing field shape but with a different phase (90^°^ phase shift). Angular deviation of PCs with respect to PC number is shown as Fig A2.D.

**Fig A2.A:**
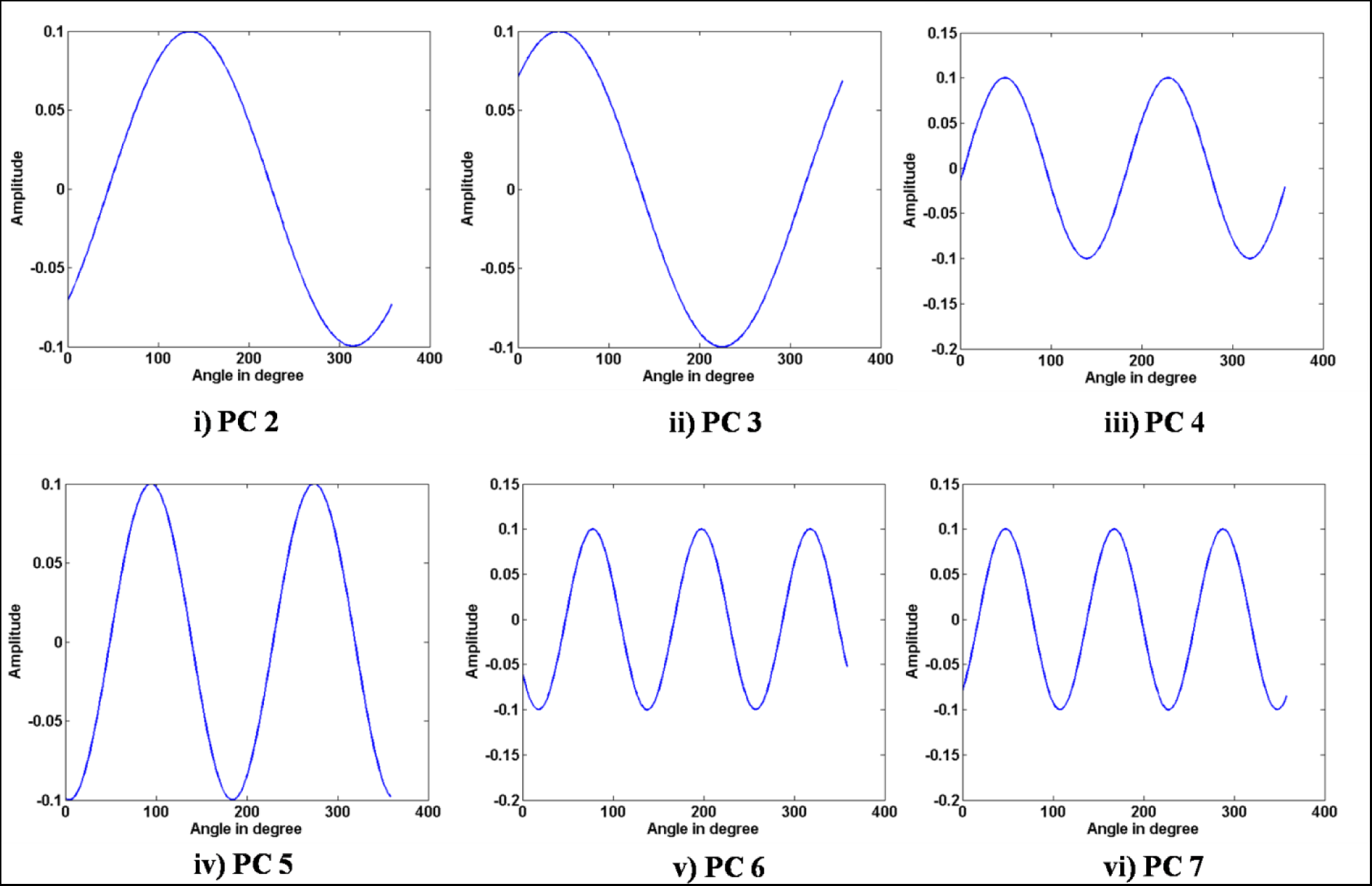
Each figure (i to vi) shows the respective sinusoidal PCs from PI layer to SC layer.

**Fig A2.B:**
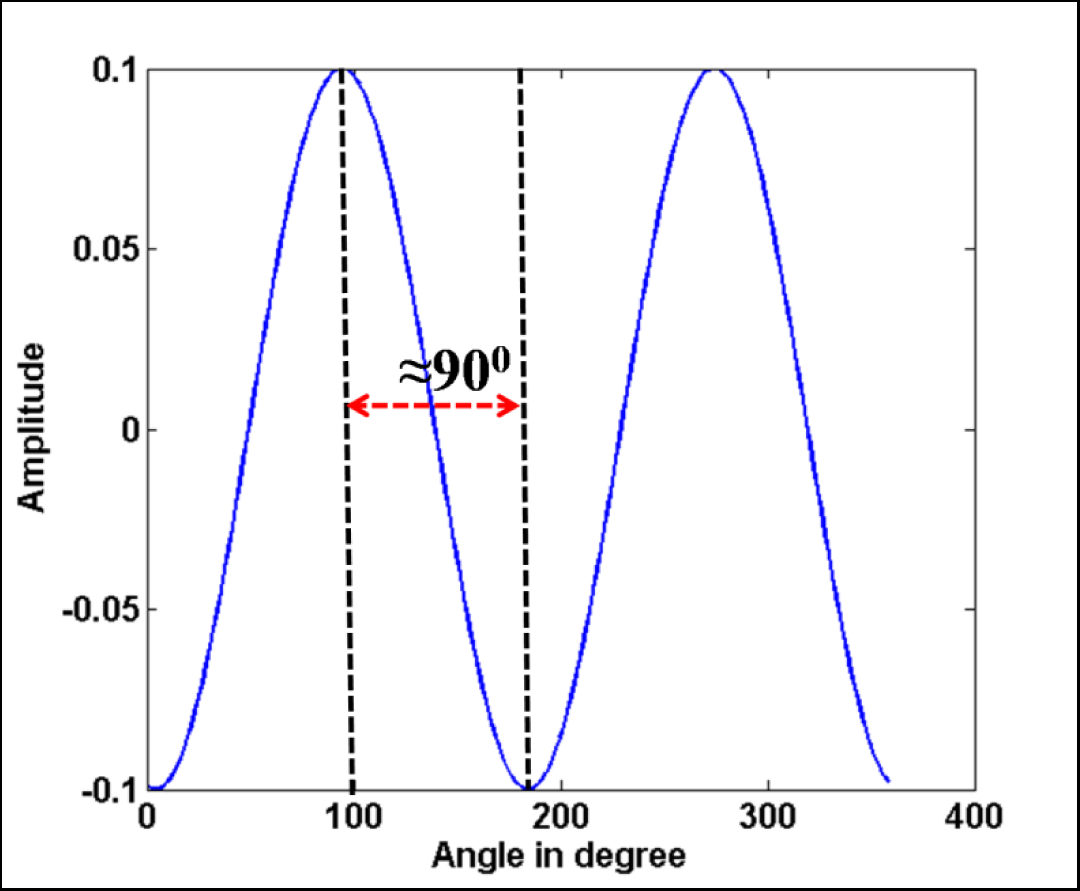
Peak to peak angular deviation of PC5. This PC gives a square shaped firing field and this is evident from the angular deviation of ≈ 90^°^ as shown in the figure.

**Fig A2.C:**
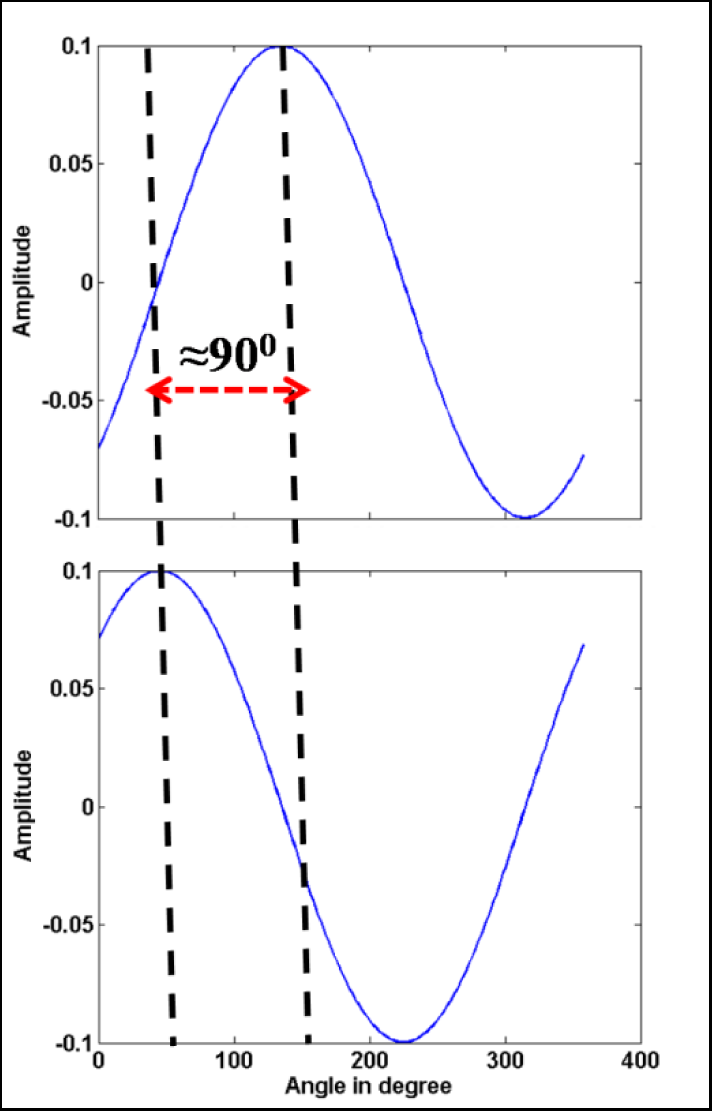
Phase offset between two pairs of PC (PC2 and PC3 shown here). This makes the respective firing fields to have a phase difference irrespective of their similarity in shape (similarity in shape is because both have the same angular deviation).

**Fig A2.D:**
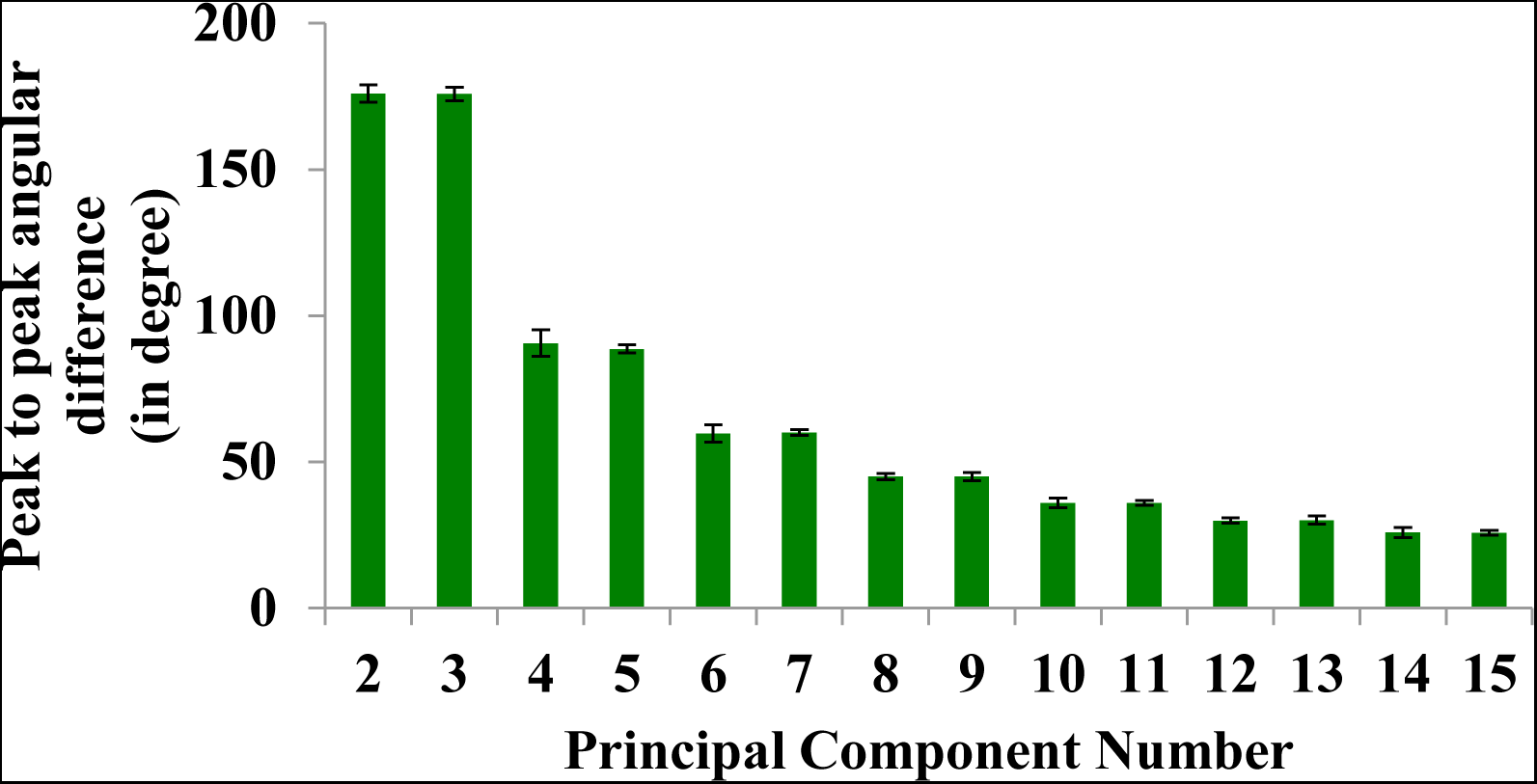
Bar graph showing the peak to peak angular deviation (y axis) of the PCs vs the respective PC number (x axis).

## A3. Modulation of limb oscillations

A virtual animal is assumed to have a rectangular strip of body with four limbs at the corners of it as shown in Fig A3.A. It is assumed to traverse a circular trajectory as shown in Fig A3.B.

**Fig A3.A:**
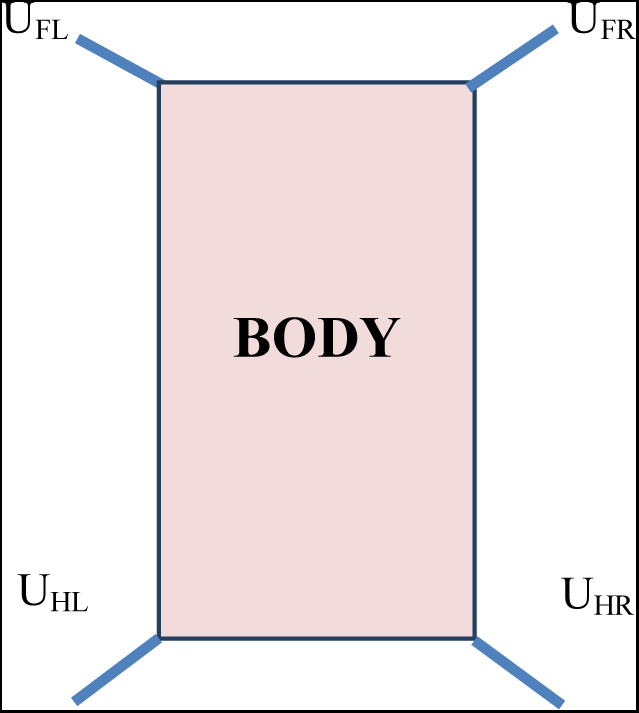
Top view of the virtual animal with rectangular body and four limbs at the corners of its body.

**Fig A3.B:**
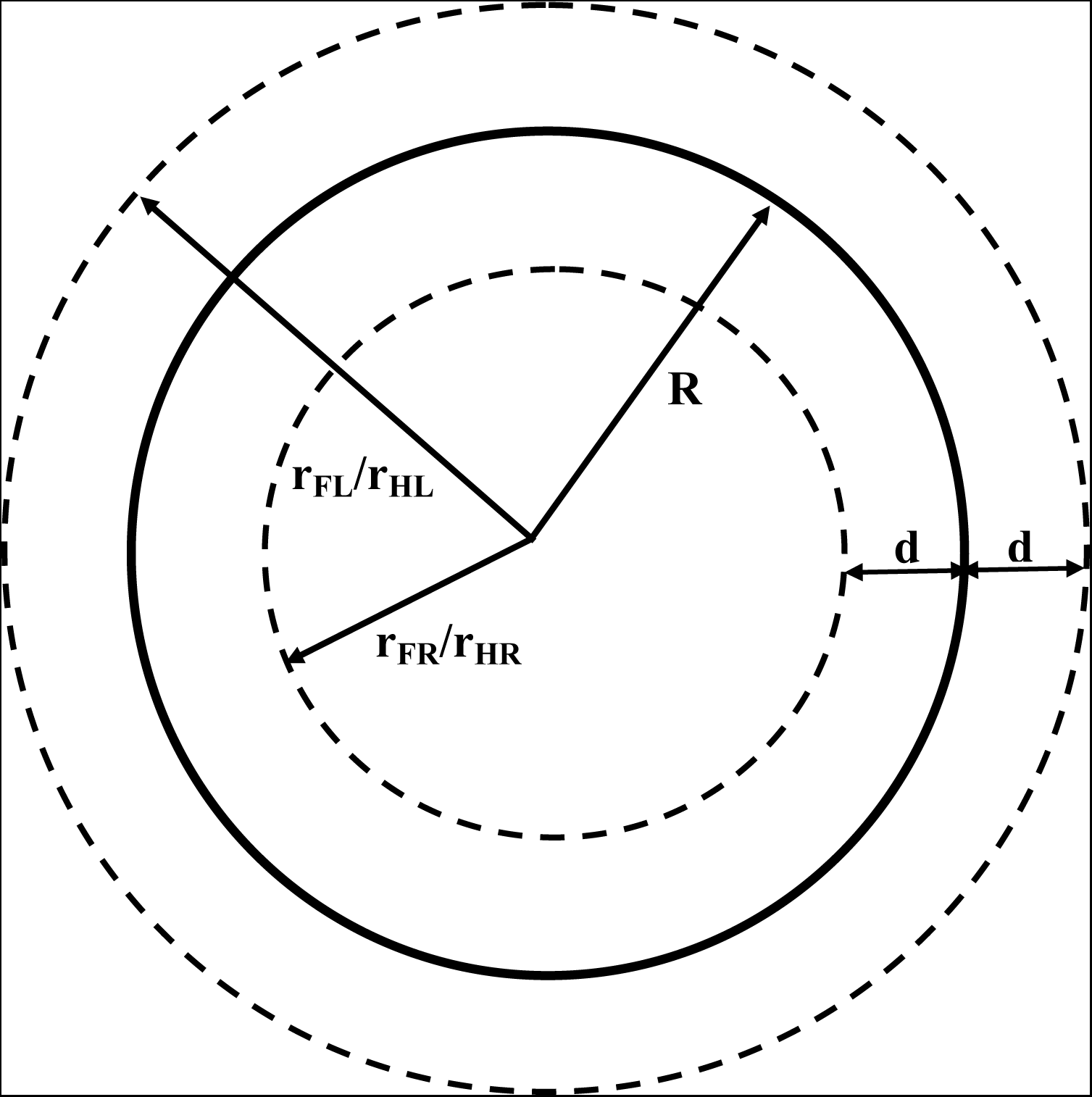
Circular trajectory traversed by the animal in anticlockwise direction. The outer dashed line indicates the trajectory traversed by both fore and hind right limbs. The inner dashed line indicates the trajectory traversed by both fore and hind left limbs. The Middle thick line indicates the trajectory traversed by the center of mass of the animal.

In Fig A3.B d is the distance between fore right and left limbs / hind right and left limbs. R is the radius of curvature of the trajectory traversed by the center of mass of the animal. r_FR_/r_HR_ is the radius of curvature of the trajectory traversed by the fore right and hind right limbs of the animal. r_FL_/r_HL_is the radius of curvature of the trajectory traversed by the fore and hind left limbs of the animal.

Each four limbs are assumed to be four oscillators. Let a_FR_ and a_FL_be the distances travelled by the fore right and left limbs respectively in one cycle of oscillation. a_FR_ and a_FL_ must satisfy the ratio given as:

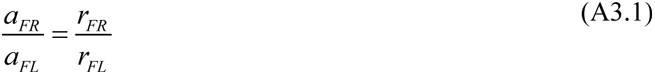

where *r*_*FR*_ and *r*_*FL*_ are given as

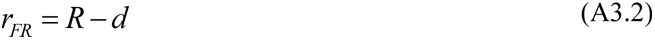

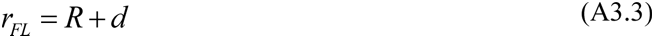

Substituting (A3.2) and (A3.3) in (A3.1),

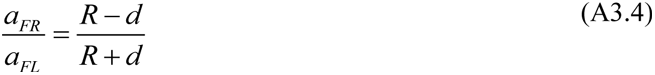

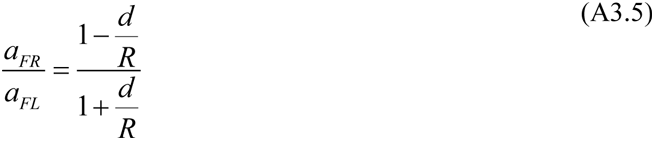

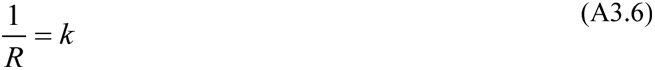

where k is the curvature of the trajectory. Substituting (A3.6) in (A3.5),

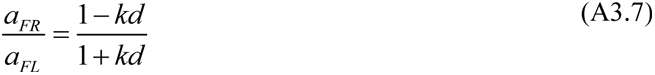

Hence,

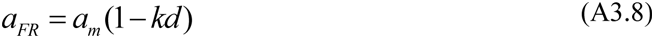

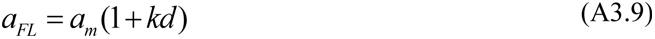

where *a*_*m*_ is the distance travelled by the center of mass in one cycle of oscillation and is given as,

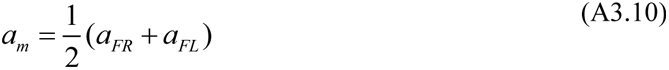

The side view of the animal is shown in Fig A3.C.

**Fig A3.C:**
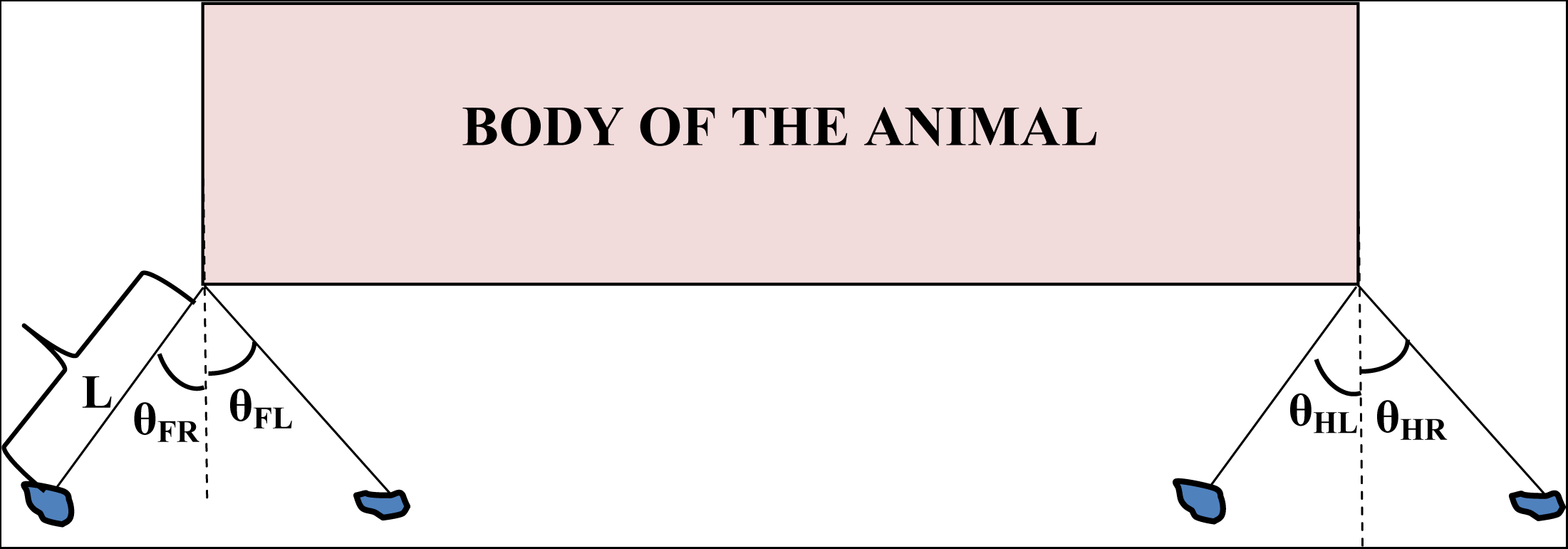
Side view of the animal. θ_FR_,θ_FL_,θ_HR_ and θ_HL_are the respective angles made by fore right, fore left, hind right and hind left limbs with the line parallel to the normal passing through the center of mass of the animal. L is the length of the animal’s leg.

In Fig A3.C the animal is assumed to navigate using trot rhythm where the contralateral limbs are in phase and ipsi lateral limbs are out of phase (Collins and Richmond, 1994) as shown in the figure. Given the length of the leg ‘L’ and the angle ‘θ’ the distance travelled by fore right and left limb during one full cycle is given as:

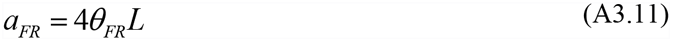

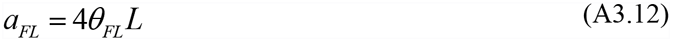

Divide (A3.11) and (A3.12). Hence we obtain

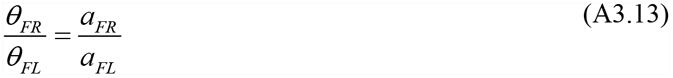

Substituting (A3.7) in (A3.13)

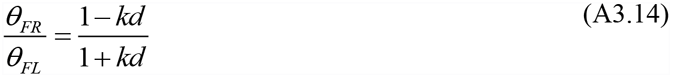

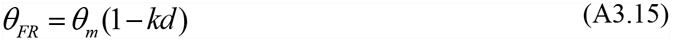

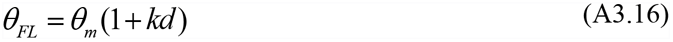

From (A3.12) a similar expression can be applied for a_m_ also such as:

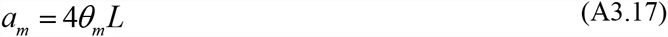

Hence,

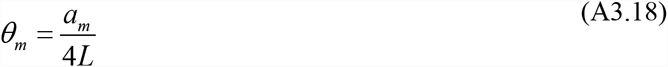

Hence the locomotor oscillatory signals from fore left and right limb are given as:

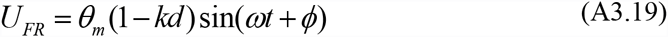

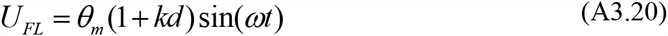

*φ* is the phase difference between the oscillations which is taken as 180^°^.
*ω* is the angular frequency of oscillations.

In the proposed model the frequency of oscillation is assumed to be a constant. It is the amplitude that varies with respect to the curvature and speed as per Eqn (A3.18,A3.19,and A3.20).

[Note: Similar Eqn. A3.1-20 hold for hind limbs too]. Hence hind limb oscillations can be expressed as:

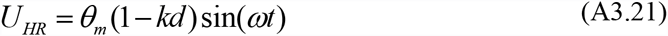

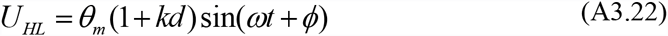

From (A3.19) and (A3.21) it is evident that

*kd*< 1. Hence there is an upper constraint on the curvature of the trajectory.

## A4. Conversion of oscillatory dynamics from Polar form to Cartesian form

Let r and θ depicts the radius and angle variable in the polar coordinates. Oscillatory dynamics in polar form can be given as

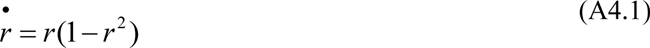

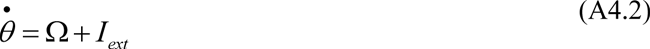

*Ω* is the base frequency of the oscillator.

I_ext_ represents the external input that affects only the phase dynamics of the oscillator.

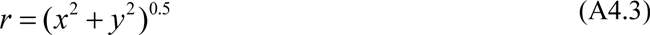

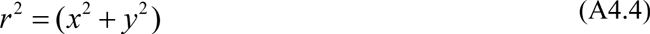

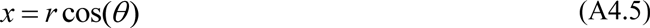

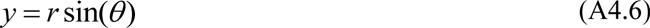

Differentiating (A4.5)

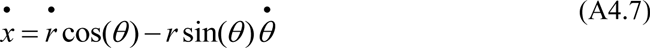

Substituting (A4.1) and (A4.2) in (A4.7)

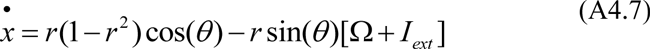

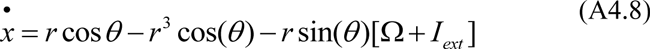

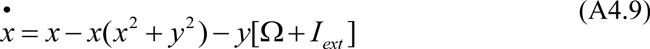

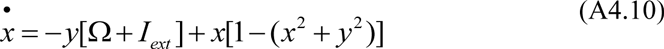

Differentiating (A4.6)

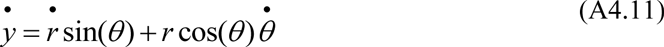

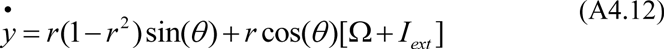

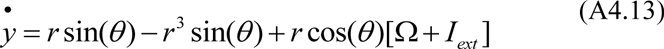

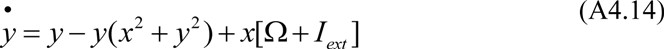

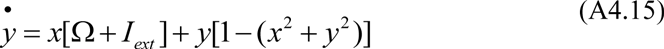

(A4.10) and (A4.15) form the dynamics of the oscillators in Cartesian form that we used in the model. From the Polar form of the dynamics (A4.1) and (A4.2) it is evident that the external input applied to the oscillator only affects the phase and not the amplitude of the oscillations.

